# Bifunctional transcriptional effector domains control gene expression pulses in an occupancy-dependent manner

**DOI:** 10.64898/2025.12.10.693551

**Authors:** Cecelia J. Andrews, Eli J. Costa, Geovanni L. Janer Carattini, Nicole V. DelRosso, Taihei Fujimori, Masaru Shimasawa, Lacramioara Bintu

## Abstract

Dynamic gene expression pulses enable adaptive response to stimuli and can be generated in natural and synthetic systems. Controlling these dynamics typically involves circuits consisting of multiple genes and transcription factors (TFs). Here, we discover a new class of bifunctional transcriptional effector domains that can first activate and subsequently repress the same gene, producing dynamic gene expression pulses from a single input. These pulse dynamics arise from distinct, temporally separated chromatin states defined by active and repressive chromatin modifications. The balance between active and repressed states is determined by the DNA occupancy of the bifunctional TF. Bifunctional domains activate at low occupancy but switch to repression at high occupancy, resulting in a non-monotonic TF input-gene expression output relationship tunable by TF concentration and number of DNA binding sites. We develop a kinetic model that links TF occupancy to gene expression transitions, allowing for the programming of eight distinct cell “states” – combinations of On/Off states of 3 reporter genes – using a single bifunctional effector. This work establishes the theoretical framework and molecular mechanisms of pulse-generating gene regulation by bifunctional domains and creates a foundation for engineering complex multi-gene circuits.

## Introduction

Transcription factors (TFs) regulate dynamic gene expression programs that allow cells to control signal responses and adaptation(*1–5*). Transient pulses of gene expression, a key feature of signal adaptation, are important in many contexts such as cell differentiation and immune responses(*6–13*). For example, Gli activity is transiently upregulated in the developing vertebrate neural tube, and the level and duration of its activity encodes cell fate decisions(*14*, *15*). These complex expression dynamics are generally thought to arise from integration of both activating and repressing inputs within gene regulatory networks(*7*, *12*, *16*, *17*). How TFs within regulatory networks control activation and repression to create dynamic expression outputs is a central question in systems biology and gene regulation.

On a molecular level, TFs control gene expression levels and dynamics through their effector domains (EDs), protein domains that recruit cofactors to regulate transcription(*18–20*). EDs are classically viewed as being either activation domains (ADs) or repression domains (RDs) that promote or inhibit transcription, respectively(*20*). However, our recent large-scale investigation of human ED functions uncovered a sizable class of bifunctional EDs that can both activate a minimal promoter and repress a constitutively active promoter(*21*) (Figure S1A). While bifunctional TFs that can activate certain genes and repress others have been reported in yeast, plant, and mammalian cells(*22–30*), control of both activation and repression through the same bifunctional ED has not been examined in detail. In particular, given the complex gene expression dynamics that arise from simultaneously activating and repressing inputs at the network level(*31*), we wondered how bifunctional domains affected gene expression over time.

Intriguingly, when individually recruited to a synthetic reporter gene, some bifunctional domains generate gene expression pulses at the level of the cell population despite constant TF input(*21*) (Figure 1A). These dynamics resemble those produced by more complex network motifs such as incoherent feed-forward loops and negative feedback(*12*, *16*). However, such network architectures are absent in this minimal synthetic system. These observations raise fundamental questions: how do individual bifunctional domains generate gene expression pulses? What parameters determine when they activate, repress, or switch between these modes? Dissecting the fundamental mechanisms and principles of this new class of human transcriptional regulators is essential for understanding how bifunctional TFs operate within the more complex endogenous gene regulatory networks involved in development and cellular signal processing. Additionally, understanding the parameters that control their behavior would enable engineering of complex transcriptional dynamics such as pulses using the smallest possible set of components, an important constraint for robust synthetic biology applications(*32–34*).

**Figure 1:**
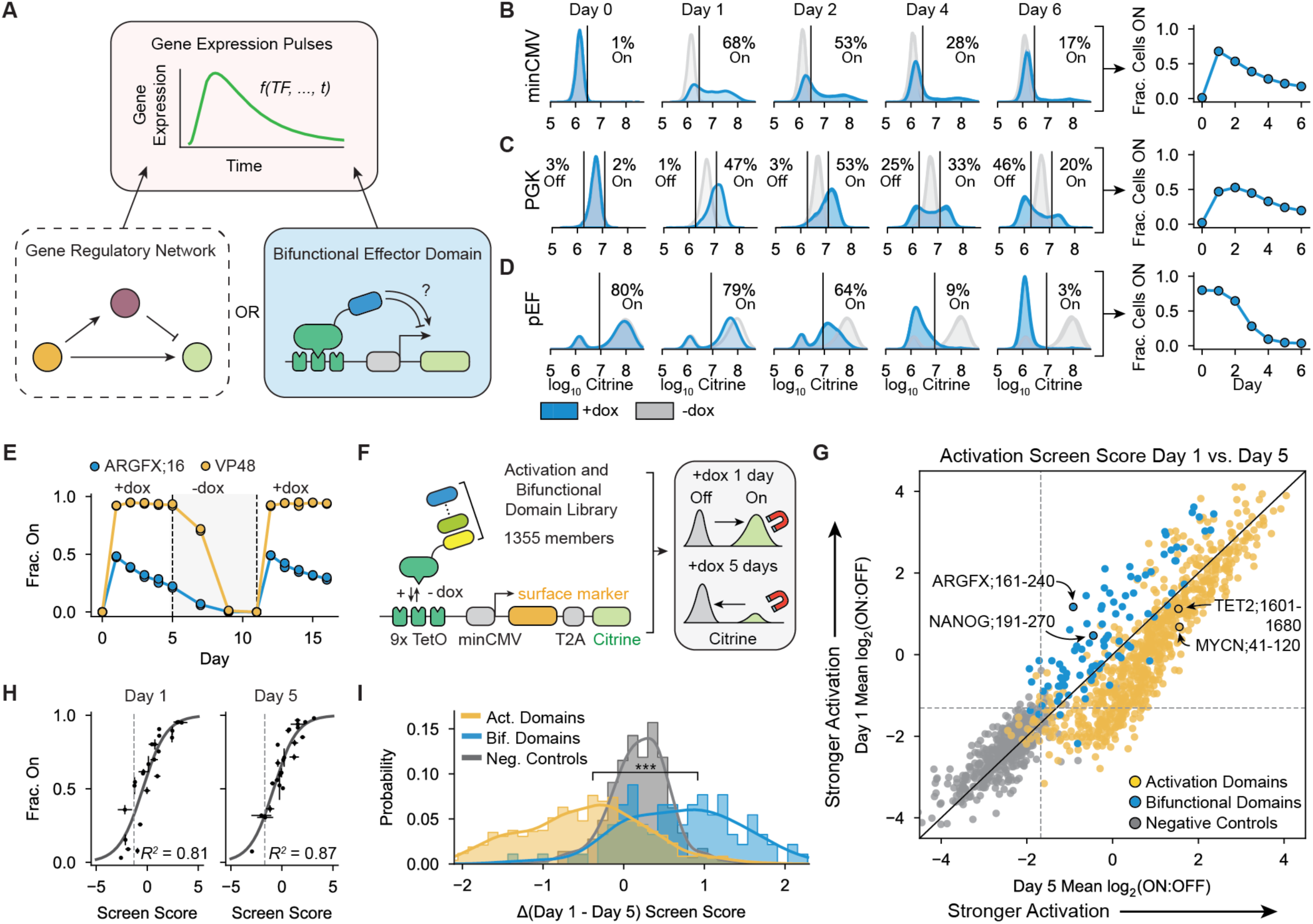
Bifunctional domains generate a pulse of gene expression at the level of the cell population when recruited to a minimal promoter. **A)** (Top box) Gene expression pulses, when expression of a target gene rapidly increases and then decreases over time, are typically thought to be controlled by gene regulatory networks (bottom left box). We find bifunctional transcriptional effector domains can also intrinsically regulate dynamic pulses of gene expression in human cells when recruited to a reporter gene (bottom right box). B-D) Left: Citrine fluorescence distributions from flow cytometry of cells where ARGFX;161-240 (ARGFX;16) was recruited to the indicated promoter for 6 days with 1000 ng/mL dox (2 biological replicates/10,000 cells/biorep, gray=no dox, blue=dox). Right: Fraction of cells with Citrine reporter active on each day of recruitment. **B)** ARGFX;16 recruited to minCMV promoter. **C)** ARGFX;16 recruited to PGK promoter. **D)** ARGFX;16 recruited to pEF promoter. **E)** Fraction of cells On during ARGFX;16 (blue) or VP48 (yellow) recruitment to the minCMV reporter from Day 0-5, followed by dox washout from Day 5 - 10 (shaded), then repeated recruitment with dox from Day 11-15. **F)** Schematic of HT-Recruit(*35*) assay to measure activation and bifunctional domain gene regulatory dynamics at minCMV. A library containing 1,355 members including activating and bifunctional effectors as well as random negative control tiles was cloned into a lentiviral vector and delivered as a pool to K562 cells containing a genome-integrated reporter (Figure S1A)(*21*). Each protein tile is expressed as a fusion to rTetR, allowing its recruitment to the reporter upon addition of 1000 ng/mL dox to the cell media. Cells were magnetically separated into on and Off fractions after 1 or 5 days of dox treatment and the domains were sequenced to determine their enrichment score in the On fraction. **G)** Average Day 1 vs. Day 5 screen score of 2 biological replicates (yellow=activation domains, blue=bifunctional domains, gray=negative controls). Dotted lines indicate activation hit threshold determined by negative control distribution. **H)** Comparison of screen scores and individually recruited tiles at minCMV. Error bars indicate the difference between 2 biological replicates. N=24 individually validated negative control, activating, and bifunctional tiles. I) Distributions of difference between average Day 1 and Day 5 screen scores for negative controls, bifunctional tiles, and activating tiles (K-S test, p<0.0001).

Here, we combine synthetic recruitment and high-throughput measurements to build a mechanistic and theoretical framework for how and when bifunctional EDs activate, repress, and produce pulses of gene expression. By profiling transcriptional dynamics for >1,000 human activation and bifunctional domains, we find that pulse generation is a general property of bifunctional EDs and is distinct from classical ADs. Mechanistically, bifunctional EDs depend on both coactivators and corepressors to establish long-lived chromatin and gene expression states, and dynamics arise from a separation of timescales between chromatin state transitions. Furthermore, we find the balance between activation and repression is controlled by TF occupancy: low occupancy favors sustained activation, whereas high occupancy drives repression and pulses, producing a non-monotonic steady-state input–output relationship tunable by TF concentration and the number of binding sites. We develop a kinetic model linking occupancy to state transitions and use it to engineer synthetic systems that create and dynamically regulate eight gene-expression states in a cell population with a single synthetic TF.

## Results

### Bifunctional domains intrinsically regulate pulses of gene expression

To determine the mechanisms and control parameters underlying bifunctional domain regulatory activity, we used a synthetic recruitment system where EDs of interest are expressed as fusion proteins with the rTetR DNA binding domain, which binds to TetO DNA sites in the presence of doxycycline (dox) (Figure S1B). These synthetic recruiter constructs are stably expressed in human K562 cells containing a genomically integrated reporter with 9 TetO sites upstream of a promoter controlling the expression of a Citrine fluorescent reporter gene and a surface marker protein (Figure S1B)(*35*). Citrine enables population-wide measurements of gene expression dynamics with flow cytometry, while the surface marker reporter allows magnetic separation for high-throughput measurements. Critically, this synthetic recruitment system is orthogonal to endogenous gene regulatory networks in human cells, allowing us to precisely quantify the gene expression dynamics regulated directly by EDs of interest without confounding factors.

We first focused on the bifunctional domain from the transcription factor ARGFX as a model for bifunctional gene regulation and pulse generation. Full-length ARGFX is an ETCHbox TF containing a single bifunctional domain(*21*) that regulates zygotic gene activation in early embryonic development(*36–38*). Notably, clusters of ARGFX-dependent genes display transient, pulse-like expression patterns(*37*). Our previous work that tested 80 amino acids (aa) consecutive fragments from human TFs identified aa 161-240 in ARGFX (denoted ARGFX;16) as a strong bifunctional effector(*21*). Under constant dox recruitment, ARGFX;16 produced clear promoter-dependent dynamics: at minCMV and PGK, most cells initially activated the reporter, then a subset of cells silenced it over time (Figure 1B-C). We observed similar gene expression dynamics when recruiting the full-length ARGFX protein (with its DBD replaced by a Glycine-Serine linker) to the minCMV promoter (Figure S1C), suggesting the short 80aa domains can recapitulate the transcriptional effector function of this TF. Unlike classical activation and repression domains, which continuously activate or repress our reporter genes, bifunctional domains perform both functions at the same gene depending on recruitment duration (Figure 1B-C). These dynamics result in a bimodal cell population after 6 days of recruitment, despite identical TF input (Figure 1B-C). In the same reporter context, ARGFX;16 strongly repressed the pEF promoter (Figure 1D). Together, these data show that bifunctional domains can drive activation, repression, or transient pulses of gene expression depending on the promoter.

Because these pulses arise in an orthogonal system without network feedback, we first asked whether they are caused by intrinsic properties of ARGFX;16 and not other cellular mechanisms. To ensure that a reduction in effector domain abundance was not the cause of the decrease in Citrine expression over time in some cells, we labeled our synthetic TF with a HaloTag during recruitment. We observed no correlation between the Citrine expression state of the cells and the normalized HaloTag ligand fluorescence, ruling out concentration changes as the cause of pulse dynamics (Figure S1D). To verify that a change in cell proliferation rate between expressing and non-expressing cells was not causing the pulse, we added ViaFluor proliferation dye to the cells at the start of recruitment. Once again, we observed no correlation between gene expression state and the intensity of the proliferation dye within the cells (Figure S1E). These results are consistent with ARGFX;16 actively regulating pulses of gene expression.

To test whether the expression pulse leaves an epigenetic memory of repression or whether the promoter resets after ARGFX;16 release, we released and re-recruited ARGFX;16 and observed a second pulse of similar magnitude upon a second dox stimulation (Figure 1E, blue line). These results indicate that ARGFX;16-mediated repression does not confer epigenetic memory and that the promoter returns to a basal state after the TF is removed. The ability to elicit repeated pulses also suggests activity-mediated transgene silencing is unlikely(*32*, *39*). Moreover, recruitment of the strong activator VP48 did not produce pulse-like dynamics (Figure 1E, yellow line), indicating that strong activation alone is insufficient for pulse generation and that the observed silencing is specific to ARGFX;16 rather than general *cis*-effects, such as squelching.

Finally, we tested the cell-type specificity of pulse-generation by ARGFX;16 by recruiting the domain to reporter genes in both hiPSCs and HEK293T cells. In both cell types, a robust pulse of gene expression was observed (Figure S1F-G). Notably, in HEK293T, where the constitutive pEF starts at a lower expression level than in K562, we observe pulsing (Figure S1G) instead of repression (Figure 1D). Altogether, these results suggest that the pulsatile dynamics observed during ARGFX;16 recruitment is caused by ARGFX;16-mediated activation, and then repression, of the reporter gene, supporting the hypothesis that pulse-generating dynamics are an intrinsic property of bifunctional domains.

### Pulse generation is a general property of human bifunctional domains

After determining that pulse-generation is an intrinsic property of the ARGFX bifunctional domain with constant input at target genes, we wondered if this property was common to all human bifunctional domains identified in our previous work screening through all human TFs and chromatin regulators(*21*). To test this hypothesis, we used HT-Recruit(*35*) to systematically measure the dynamic gene regulatory functions of a library containing 1,355 previously annotated activating and bifunctional effector domains from human TFs(*21*) (Figure 1F, Table S1). Domains were classified as bifunctional by their ability to activate the minimal minCMV promoter after 2 days of recruitment and repress the constitutively active promoter pEF after 5 days of recruitment in previous work, or as activation domains if they only activated the minimal promoter.

To measure the function of each ED over time in our pooled assay, we recruited each library member to minCMV by adding dox, then magnetically separated On and Off cells after 1 day and 5 days (Figure 1F). We calculated an enrichment score in the On fraction for each ED by sequencing (Methods, Figure 1G, Figure S1H-I). We found 84% (76/91) of bifunctional domains had a higher enrichment score in the On fraction after 1 day of recruitment compared to 5 days of recruitment, suggesting they are pulse-generating: they initially activate the reporter gene before switching to repression (Figure 1G, Table S1). In contrast, 84% (606/724) of activation domains had the same or higher enrichment score after 5 days of recruitment (Figure 1G, Table S1), suggesting they continue functioning as activators at minCMV and increase gene expression over time.

We validated the dynamics measured in our HT-recruit assay through individual flow cytometry experiments for 24 domains (Figure 1H, S1J, Table S2), and observed strong agreement between the fractions of cells expressing Citrine after 1 and 5 days of recruitment and the corresponding screen scores for both day 1 and day 5 (Figure 1H). This agreement indicates our screening strategy can accurately assess both the strength of initial activation and subsequent repression of the minCMV promoter and therefore measure gene expression dynamics for each domain. We quantified the shape of the pulse using 2 key shape measurements: Amax, the amplitude of the pulse, and the magnitude of relative repression Δ, as the difference between Amax and the fraction of cells expressing Citrine at the end of recruitment (Figure S1K). Individually validated bifunctional domains sampled a range of Amax and Δ values (Figure S1L), showing the pulse shape varies depending on domain identity.

By comparing the difference in screen scores between day 1 and day 5, we observe a clear separation between the dynamics of activation domains and bifunctional domains at the minCMV reporter, with almost all bifunctional domains having a positive difference (less activation on day 5) and almost all activation domains having a negative difference (more activation on day 5) (Figure 1I). Altogether, our high-throughput screen supports the conclusion that pulse generation is a shared property of human bifunctional domains, potentially enabled by their unique ability to also repress transcription which is not shared by canonical activation domains.

### A minimal kinetic model of pulse-generation by bifunctional domains

After confirming that pulse-generation is an intrinsic and shared property of most tested bifunctional domains, we sought to develop a minimal model that could explain these gene expression dynamics and make quantitative, testable predictions. Given that these domains can both activate and repress transcription and create bimodal Citrine expression distributions during individual recruitment assays, we reasoned that the observed dynamics are consistent with a three-state kinetic model of gene expression states (Figure S1M, Figure 2A). In this model, recruitment of a bifunctional domain causes cells to stochastically transition from a “ground” gene expression state (G, set by the promoter) to an “active” state with high Citrine expression (A) and a “silent” state with no expression (S). All three states are evident during recruitment to the PGK promoter, where most cells first increase in Citrine expression above the baseline level and then a fraction become silenced to Citrine levels below baseline (Figure 1C, Figure S3A). Based on our observation that the system can re-pulse after dox release (Figure 1E), we assume all cells return to the ground state in the absence of recruitment. We assume that cells in both the ground and active states can be silenced by a dominant mechanism of repression, as was previously observed for many combinations of activation and repression domains(*40*). Finally, we assume that bifunctional domain recruitment promotes transitions to the active state at rate *k_1_*and to the silent state at rate *k_2_*. These assumptions are sufficient to predict a pulse of gene expression in the cell population (Figure 2B, black line) using just these two parameters.

**Figure 2:**
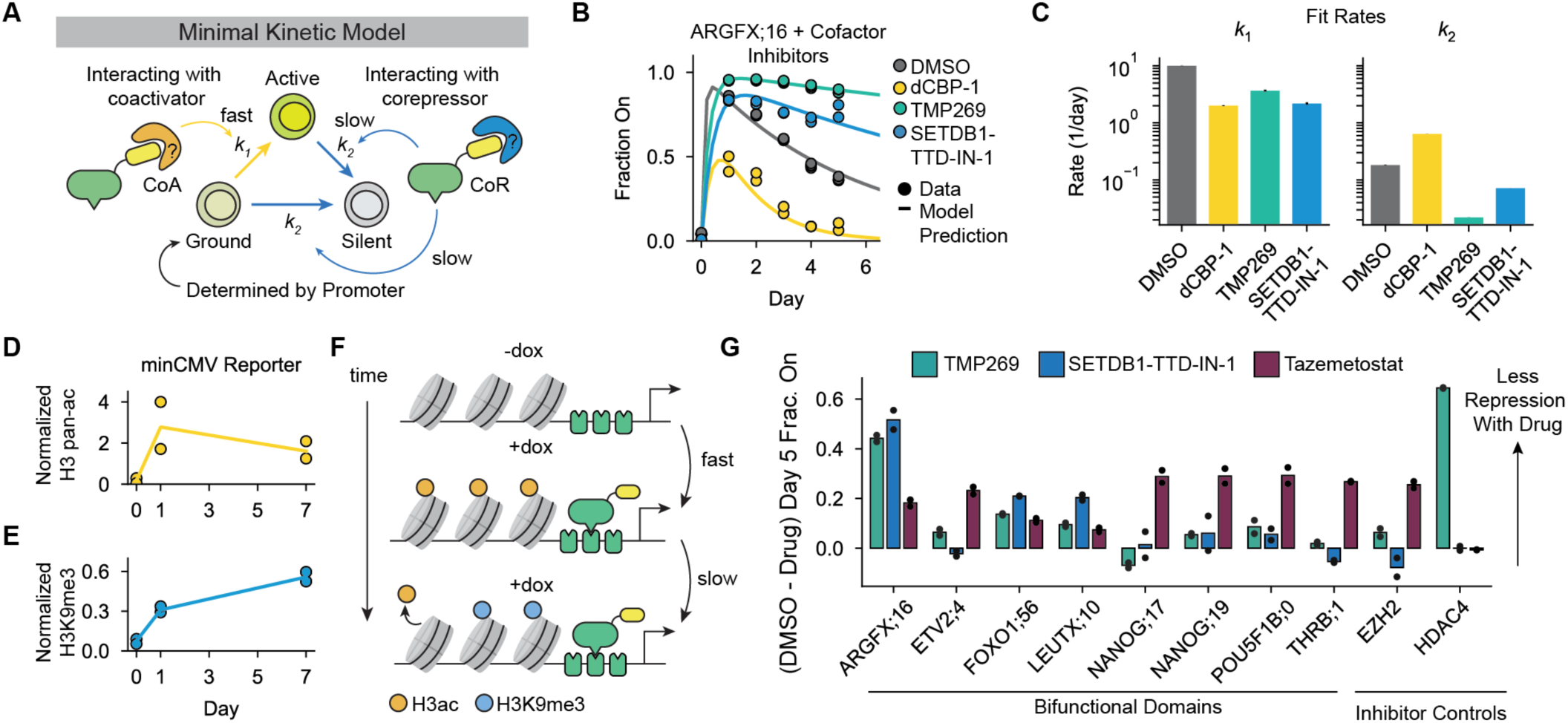
Pulse-generating regulatory dynamics depend on chromatin regulators that produce distinct chromatin states over time. **A)** Schematic of minimal 3-state kinetic model. The activation rate (*k_1_*, yellow arrow) is regulated by the bifunctional ED (yellow) interaction with transcriptional coactivators (orange). The silencing rates (*k_2_*, blue arrows) are regulated by interaction with transcriptional corepressors (blue). **B)** Fraction of cells with Citrine reporter On during ARGFX;16 recruitment to minCMV in media containing 1000 ng/mL dox and 10 μM dCBP-1 (P300/CBP PROTAC degrader, yellow), 10 μM TMP269 (Class IIa HDAC inhibitor, green), or 10 μM SETDB1-TTD-IN-1 (SETDB1 inhibitor, blue) or DMSO (vehicle control, gray). (dots=data, lines=model fit to data in given inhibitor condition). **C)** Fit rates *k_1_* (activation rate) and *k_2_* (repression rate) in each inhibitor condition based on the data from (B). **D)** Normalized CUT&RUN H3 pan-acetylation reads/kb for the minCMV reporter gene after 0, 1, or 7 days of ARGFX;16 recruitment with 1000ng/mL dox. Data are normalized to GAPDH (H3 pan-acetylation positive control gene) and ZNF140 (H3 pan-acetylation negative control gene). **E)** Normalized CUT&RUN H3K9me3 reads/kb for the minCMV reporter gene after 0, 1, or 7 days of ARGFX;16 recruitment with 1000ng/mL dox. Data normalized to ZNF140 (H3K9me3 positive control gene) and GAPDH (H3K9me3 negative control gene). **F)** Schematic showing that H3 acetylation reaches maximum levels at the reporter faster than H3K9me3. **G)** Difference in fraction of cells with Citrine On after 5 days of recruitment with 1000ng/mL dox in either 10 μM TMP269 (green), 10 μM SETDB1-TTD-IN1 (blue), or 10 μM Tazemetostat (EZH2 inhibitor, maroon) compared to DMSO for the different bifunctional tiles indicated on the x-axis. EZH2 is a positive control for inhibition by Tazemetostat and HDAC4 is a positive control for inhibition by TMP269.

By regulating both activation (*k_1_)* and repression (*k_2_)* in this 3-state kinetic model, bifunctional domains can generate transient pulses of gene expression of various shapes. We considered how these parameters affect the shape measurements Amax and Δ (Figure S1K). Amax is greatest when *k_1_* is high and *k_2_* is close to 0, and the greatest Δ occurs when *k_2_* is relatively high but *k_1_* > *k*_2_ (Figure S1N, contours). We analyzed this type of variation in bifunctional domain dynamics at minCMV by individually recruiting 11 bifunctional tiles from 9 unique TFs for 5 days (Table S2). By fitting our minimal model for each tile, we examined the variance in their rates of activation (*k_1_*), silencing (*k_2_*), and their observed shape parameters, Amax and Δ (Figure S1N). The estimated *k_1_* and *k_2_* each spanned 2 orders of magnitude and the pulse shapes could be explained by the minimal model, suggesting that bifunctional domains vary in their activating and repressing dynamics at minCMV to create pulses with differing amplitudes.

### Bifunctional domains generate a pulse of expression through recruitment of both coactivators and corepressors

Previous studies of bifunctional TFs have demonstrated certain TFs can interact with either transcriptional machinery or chromatin regulators in a context-dependent manner(*41*, *42*). Alternatively, recent studies have proposed theoretical models where TFs could regulate different kinetic steps of transcription, resulting in a loss of activation under certain conditions(*43*, *44*). Given that pulses of gene expression occur through transitions between discrete Citrine expression states on the timescale of days, we hypothesized that bifunctional domains could affect activation and repression mechanisms by recruiting chromatin regulators to promote or inhibit gene expression (Figure 2A).

To investigate the mechanisms underlying pulse-generating regulation, we tested the effects of transcriptional cofactor perturbations on the dynamics of ARGFX;16-controlled reporter expression. Full-length ARGFX has previously been shown to interact with both activating and repressing chromatin regulating complexes, including major coactivator histone acetyl transferases P300/CBP and the histone deacetylase (HDAC)-containing HES1 corepressor complex(*45*). We first tested ARGFX;16’s dependence on P300/CBP by degrading both proteins with the dCBP-1 PROTAC(*46*). P300 and CBP deposit histone 3 acetylation (H3ac), including H3K27ac that marks active promoters and enhancers(*47–49*). P300/CBP degradation partially reduced the maximum fraction of active cells (Amax) but did not reduce the relative repression, Δ (Max - Day 5), compared to DMSO, suggesting histone acetylation is important for ARGFX;16-mediated activation but not repression (Figure 2B, yellow; Figure S2A-B). Small molecule inhibition of P300/CBP acetyltransferase activity(*50*) also reduced Amax(Figure S2C).

Given histone acetylation is required for activation, we hypothesized that removal of acetylation by histone deacetylases (HDACs) would be required for repression. We found Class IIa HDAC inhibition with TMP269(*51*) completely abolished repression at minCMV without affecting activation, such that Amax was unchanged but Δ(Max - Day 5) was greatly reduced (Figure 2B, green). HDAC inhibition also reduced repression of the pEF promoter (Figure S2D-F). We further tested the necessity of other chromatin regulators involved in major transcriptional repression pathways: SETDB1, the writer of H3K9me3, and EZH2, the writer of H3K27me3. We found inhibition of SETDB1 with SETDB1-TTD-IN-1 resulted in complete loss of repression (Figure 2B, blue), while inhibition of EZH2 had a minimal effect on repression at the minCMV and pEF promoters (Figure S2G-J), indicating SETDB1 and H3K9me3 are required for repression.

We fit our minimal kinetic model (Figure 2A) to our cofactor inhibitor data and found P300/CBP degradation reduced the activation rate *k_1_* while HDAC inhibition reduced the silencing rate *k_2_* (Figure 2B-C). This suggests activation (*k_1_*) is related to the rate of histone acetylation while silencing (*k*_2_) is related to the rate of histone deacetylation and H3K9 methylation. Furthermore, *k_1_* is much larger than *k_2_*, indicating the mechanism of activation affects gene expression more quickly than the mechanism of repression. To determine the dynamics of histone acetylation, deacetylation and methylation, we performed CUT&RUN for H3 pan-acetylation and H3K9me3 after 0, 1, or 7 days of recruitment. Before recruitment (day 0), our minCMV reporter does not have significant levels of acetylation or methylation compared to endogenous control genes (Figure 2D-E, S2L-M, Methods). After 1 day of ARGFX;16 recruitment, our bulk measurements show both histone acetylation and H3K9me3 present at the locus at significant levels above no dox conditions (Figure 2D-E, S2L-M Day 1), even though at this timepoint 79% cells were in the On state (Figure S2K). This could arise from separate molecules having either H3ac or H3K9me3, bivalent single molecules, or a combination of both. After 7 days of recruitment, histone acetylation at the locus was similar or slightly lower compared to day 1, while H3K9me3 was increased (Figure 2D-E, S2L-M, Day 7), corresponding with an increase in percentage of cells Off from day 1 to day 7 from 21% to 82% (Figure S2K). Consistent with our model predictions, H3 pan-acetylation reached its maximum after 1 day of recruitment, while H3K9me3 levels continued increasing over 7 days of recruitment. These observations are consistent with measured rates of histone acetylation and deacetylation, which occur on the order of minutes(*52*), compared to H3K9me3, which propagates on the order of hours to days(*53*). These data suggest recruitment of both coactivators and corepressors to the reporter gene results in temporally separated chromatin state transitions (Figure 2F), leading to a pulse of gene expression.

We additionally wondered whether bifunctional domains from different TFs in our study shared a common mechanism of repression or if mechanisms differed between domains. We recruited bifunctional domains from 7 different TFs to the minCMV reporter while inhibiting major chromatin repressor complexes and compared the difference in Day 5 fraction On to the DMSO control condition. We found bifunctional domains varied in their dependence on EZH2, SETDB1, and Class IIa HDACs to repress minCMV after initial activation (Figure 2G). Only 2 other domains resembled the ARGFX;16 response, being more sensitive to HDAC and SETDB1 inhibition compared to EZH2 (Figure 2G, domains from FOXO1 and LEUTX). The remaining 5 of 8 domains tested were most sensitive to EZH2 inhibition (Figure S2G, domains from ETV2, NANOG, POU5F1B and THRB). Bifunctional domains can therefore use different mechanisms of repression to control the pulse behavior.

### Long-term recruitment of bifunctional domains creates stable activated and silenced gene expression states

We noticed that the ARGFX bifunctional domain was able to silence almost all cells when recruited to the constitutively active pEF promoter (Figure 1D). We wondered if the recruited corepressors would completely dominate and silence all cells during longer recruitment to the minCMV promoter, as predicted by our simple three-state model (Figure 2A). To test this prediction, we recruited the ARGFX bifunctional domain upstream of the minCMV promoter continuously for 15 days. Interestingly, we found that the cell population reached a steady state after 10 days of recruitment with ∼21% of cells still expressing Citrine (Figure 3A). We reasoned that a non-zero steady state of active cells could either be achieved through a dynamic equilibrium between cells in active and silent gene expression states (Figure 3B, Dynamic Equilibrium), or by cells stochastically committing to stably active and silent states over time (Figure 3B, Divergent States). Both models reasonably fit the gene expression dynamics during ARGFX;16 recruitment to the minimal promoter and can accurately describe the non-zero steady state fraction of active cells (Figure S3B-E).

**Figure 3:**
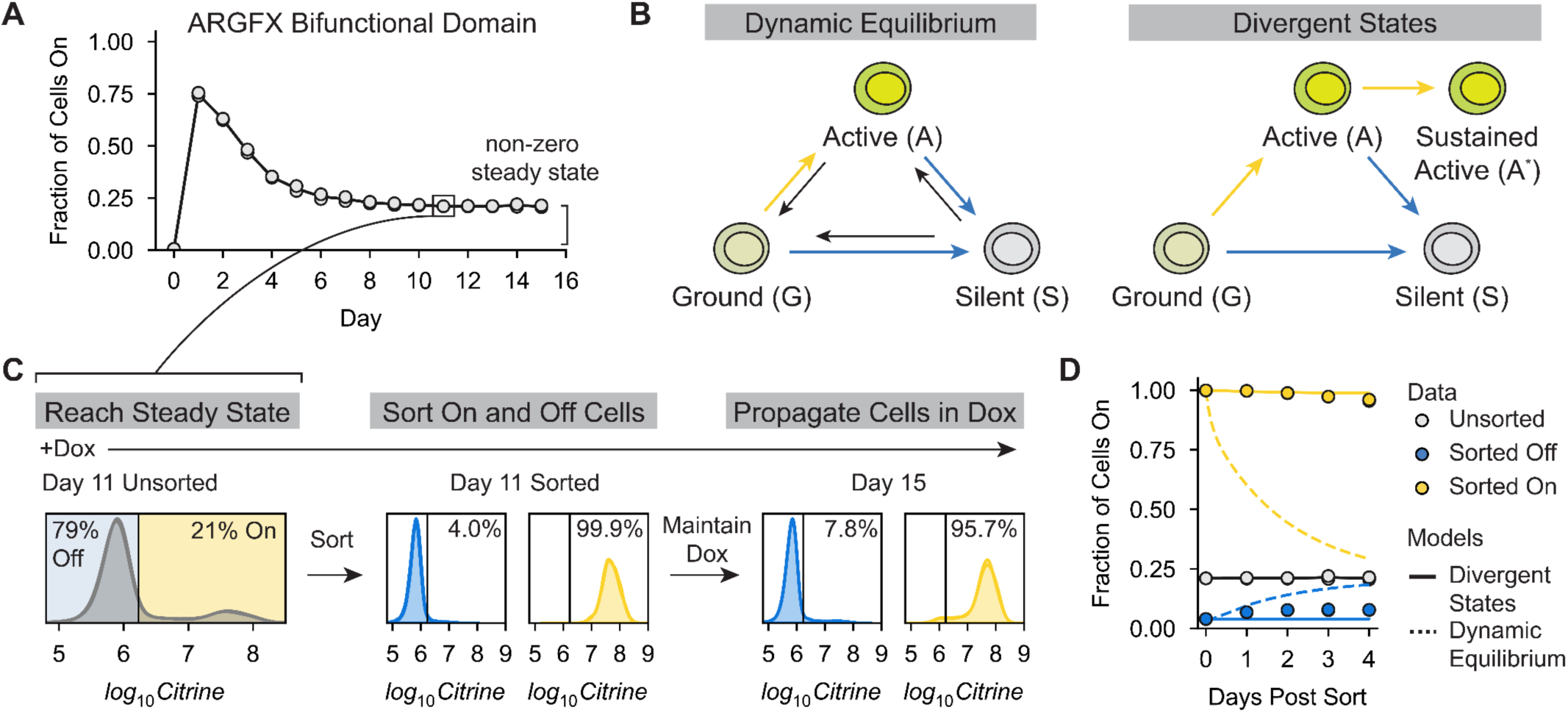
Long-term continuous recruitment of a bifunctional domain produces stable active and silenced gene expression states. **A)** Fraction of cells expressing Citrine during ARGFX;16 recruitment to minCMV for 15 days. The time of sorting is indicated with a black box. **B)** Models that can reproduce the observed non-zero steady state fraction of active cells observed during ARGFX;16 recruitment. Yellow arrows indicate transitions to Citrine On states, blue arrows indicate transitions to Citrine Off states. Black arrows represent back rates needed for a dynamic equilibrium. Cells in the ground state are assumed to not express Citrine. **C)** Sorting procedure to distinguish the possible models in B. (Left) Citrine fluorescence distribution on Day 11 of ARGFX;16 recruitment at minCMV with 1000ng/ml dox. (Middle) Citrine fluorescence distributions of cells sorted Off (blue) and On (yellow) the day of sorting. (Right) Citrine fluorescence distributions of sorted cell populations on Day 15 of recruitment, 4 days after sorting. Cells are kept in dox throughout the experiment. **D)** Fraction of cells expressing Citrine over time for unsorted (gray), sorted On (yellow), and sorted Off (blue) populations. Dashed lines show predictions of the dynamic equilibrium model (B, left) for sorted On (yellow) and Off (blue) cell populations, solid lines show prediction of the divergent states model (B, right). Model fits used to predict gene expression dynamics are shown in Figure S3B-E.

To distinguish between these possible models, we recruited the ARGFX bifunctional domain to minCMV for 11 days, enough time for the system to reach steady state, and then sorted active and silent cells into separate populations while maintaining dox (and therefore recruitment) at all times. We then propagated each population for 4 more days in dox (Figure 3C). By sorting the cells, we altered the relative abundance of each gene expression state in the population. The dynamic equilibrium model, fit to the gene expression dynamics pre-sorting, predicts that each sorted population will reach the same steady state fraction of active cells as the initial unsorted population with roughly the same kinetics (Figure 3D, dashed lines; Figure S3C). In contrast, the divergent state model fit to the same pre-sort data predicts no re-equilibration between the sorted populations (Figure 3D, solid lines; Figure S3E). We observed negligible re-equilibration between the active and silent states over 4 more days of ARGFX;16 recruitment, as predicted by the divergent states model (Figure 3D). Therefore, we conclude that long term recruitment of ARGFX;16 causes the formation of both active and silent gene expression states that are stable while the domain is recruited at the reporter, with some cells passing through an intermediate, transiently active state before stochastically committing to sustained activation or silencing.

### Bifunctional domain occupancy determines pulse amplitude and dynamics

Cells typically respond to changes in signaling or external stimuli through cascades that culminate in changes in the nuclear abundance of TFs. How changes in TF abundance are translated into changes in gene expression can vary between systems(*54*, *55*). Because bifunctional domains encode unique gene expression dynamics and recruit both coactivators and corepressors, we wondered how changes in TF abundance would be translated into changes in the rates of activation, repression, and the resulting gene expression dynamics. Using our synthetic expression system, we can continuously and directly tune effective TF abundance by changing the dox concentration used to recruit the bifunctional domains to our reporter gene, as dox changes the amount of rTetR available to bind at the TetO sites.

To test the dose dependence of bifunctional domains, we first recruited ARGFX;16 with dox concentrations in the culture media varying between 0-1000 ng/mL and measured Citrine expression dynamics over 5 days. Surprisingly, we observed that at low dox concentrations (10-100ng/mL) the ARGFX bifunctional domain acted only as a strong activator (Figure 4A; Figure S4A, left). The fraction of active cells reached its maximum at very low dox concentrations (25 ng/mL), and this maximum was mostly consistent between 25-1000 ng/mL dox, with a slight decrease in amplitude between 500-1000 ng/mL dox. Past a concentration threshold of 100ng/mL dox, ARGFX;16 generated a pulse of gene expression with the amount of repression increasing continuously with dox (Figure 4A; Figure S4A, left). A similar concordance between dox concentration and the strength of silencing by ARGFX;16 was observed at the pEF and PGK promoters (Figure S4B-D). In contrast to the ARGFX bifunctional domain, gene expression control by the strong activator VP48 displayed minimal silencing at high dox and the strength of activation increased monotonically with domain abundance, as expected (Figure S4E, left). These data reveal that dynamics of ARGFX;16-mediated activation and repression have different sensitivities to domain abundance. The maximum fraction of active cells is very sensitive with respect to dox concentration, setting the threshold for activation at very low input concentrations, while the rate of silencing is low-pass filtered: the rate of silencing is close to 0 at lower dox, then enters a regime where the rate of silencing increases gradually with TF abundance.

**Figure 4:**
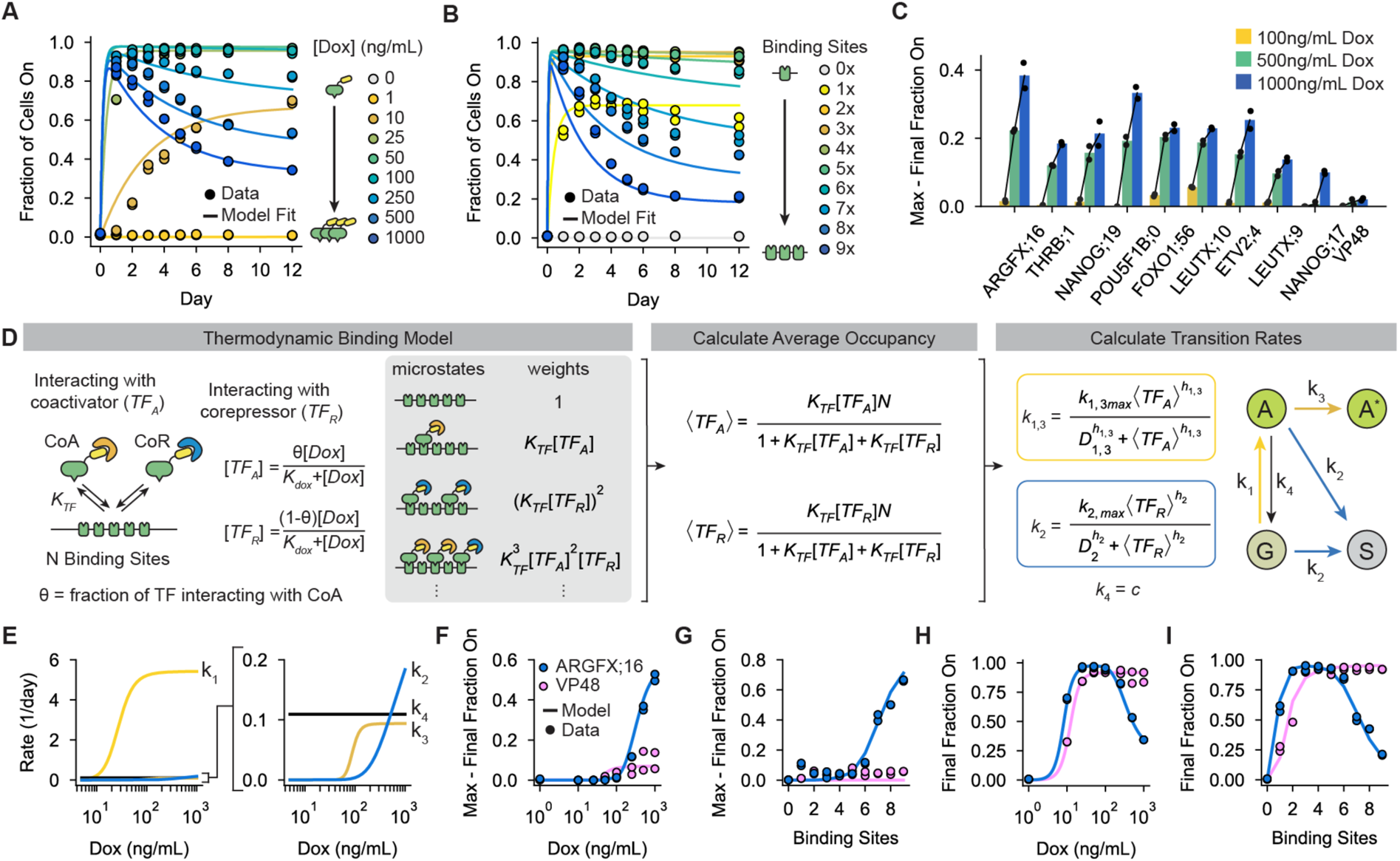
Bifunctional domain occupancy on regulatory DNA controls their relative activation and repression strengths. **A)** Fraction of cells expressing Citrine during ARGFX;16 recruitment to minCMV with varying dox concentrations, 9 TetO sites. Dots indicate measurements, lines indicate model predictions (D-I). **B)** Fraction of cells expressing Citrine during ARGFX;16 recruitment to minCMV with varying numbers of TetO sites in the reporter gene, 1000ng/mL dox. Dots indicate measurements, lines indicate model predictions (D-I). **C)** Difference between the maximum fraction of cells expressing Citrine and the final fraction of cells expressing Citrine for 9 bifunctional domains recruited for 5 days to minCMV using 3 dox concentrations. VP48 is shown as an activator-only control. **D)** Integrated model that uses a thermodynamic binding model (left) to predict TF occupancy (middle), which then serves as inputs to the kinetic, state-based model (right) developed in Figure 3B. *K_TF_* is the association constant between TetO sites and *TF_A_* and *TF_R_*, *θ* is the fraction of total TF interacting with the coactivator, *K_dox_* is the dissociation constant between rTetR and molecules of dox, and *n_A_* and *n_R_* are the numbers of activating and repressing TF bound respectively on individual DNA molecules. Rates towards Citrine On states (yellow, *k_1_, k_3_*) depend on average occupancy of the activating form〈TF_A_〉while rates towards Citrine Off states (blue, *k_2_*) depend on average occupancy of the repressive form〈TF_R_〉. *k_4_* (black) is assumed to be occupancy independent. G = ground state, A = active state, S = silent state, A* = sustained active state. Maximum transition rates are allowed to be free parameters when fitting to each dataset, all other parameters are fit globally. A full description of the model is in the Supplementary Text. **E)** Extracted transition rates from fitting the model in D to the data in A and B (Figure S4I). Maximum transition rates are taken from fitting to the dox titration dataset in A. **F, G)** Difference between the maximum fraction of cells expressing Citrine and the final fraction of cells expressing Citrine vs dox concentration (F) and number of binding sites (G) for ARGFX;16 (blue) and VP48 (pink) at minCMV (dots=data, line=model prediction). **H, I)** Final fraction of cells expressing Citrine vs dox concentration (H) and number of binding sites (I) for ARGFX;16 (blue) and VP48 (pink) at minCMV. Dots represent measurements and solid lines represent model predictions (Figure S4I,J). VP48 data were fit to a model where the rate of silencing (blue, *k_2_*) was assumed to be occupancy independent.

Because changing the dox concentration changes the amount of binding to TetO sites(*56*), we wondered if changes in domain occupancy at the regulatory region of our reporter were responsible for the observed concentration dependencies of activation and repression. To test this hypothesis, we sequentially removed TetO sites in our reporter, recruited ARGFX;16 at high dox (1000ng/mL), and measured Citrine expression over time using flow cytometry. Consistent with our previous results, at low numbers of binding sites (1-5 sites) the ARGFX bifunctional domain only activated the reporter over 12 days of recruitment (Figure 4B). As the number of binding sites increased further (6-9 sites), ARGFX;16 began to generate pulses of gene expression with the strength of silencing increasing with the number binding sites (Figure 4B; Figure S4A, right). For the VP48 activator control, we observed a monotonically increasing relationship between the strength of activation and the number of binding sites with no silencing at any number of TetO (Figure S4E, right).

Together, these results suggest that the balance between activation and repression for the ARGFX bifunctional domain is controlled by domain occupancy at the regulatory region of the reporter. Low domain occupancy favors activation, while high occupancy allows for repression. To test whether this occupancy-dependence is general among bifunctional domains, we recruited 8 other pulse-generating bifunctional domains to our minimal promoter for 5 days with lower dox concentrations (Figure S4F). The amount of silencing increased with dox concentration for each domain tested, as observed for ARGFX;16 (Figure 4C). This suggests that the activation and repression strengths of other bifunctional domains have similar occupancy dependencies, and that this principle of bifunctional gene regulation may be general across domains.

### An integrated thermodynamic and state-based model accurately describes the occupancy dependence of gene expression pulses

Our data have shown that both *cis* (i.e. binding site number) and *trans* (i.e. TF concentration) inputs establish bifunctional domain occupancy at regulatory regions, which in turn determines the relative strengths of activation and repression at target genes and the resulting gene expression dynamics. We sought to expand our previous model of bifunctional gene regulation to incorporate these two distinct inputs and gain a quantitative, predictive understanding of when a bifunctional domain will act as an activator, repressor, or pulse-generator (Figure 4D).

In order to connect changes in reporter configuration and TF concentration with transition rates between gene expression states, we developed a thermodynamic binding model of bifunctional domains where coactivator-interacting and corepressor-interacting domains compete for a fixed number of binding sites (Figure 4D, left). Because the necessary sequences for ARGFX-mediated activation and repression both map to the same small (14 residue) sequence(*21*), we did not include domains interacting with both coactivators and corepressors simultaneously in our model. By enumerating all molecular states and their respective energies, we obtain the partition function of the system and can calculate the average number of activating and repressing forms of the domain bound at the reporter for a given TF concentration and number of binding sites (Figure 4D, middle). We then assume that transition rates between gene states (active, silent, ground, etc.) can be modeled as Hill functions, with the average occupancy of the TF activating form determining the transition rates to active states and the average occupancy of the TF repressing form determining transition rates to the silent state (Figure 4D, right; Figure S4G).

We then fit this model to our dox titration and binding site perturbation data. While the simple divergent states model introduced previously (Figure 3B, right)—with added TF occupancy— is successful in describing gene expression dynamics at high binding site numbers and dox concentrations, it cannot fit the data at low binding site numbers and dox concentration (Figure S4H). To account for the low fraction of active cells at very low dox concentrations and TetO numbers, we introduced a transition from the transiently active to the ground state (*k*_4_, Figure 4D, right). We assumed this rate to be occupancy independent, and we estimate it to have a low magnitude compared to the maximal rate of activation (*k*_1_) based on fits to experimental data (Figure S4I). With this updated model, all parameters were globally fit between the two datasets (varying dox and varying TetO number) except for the maximum values of the transition rates, which we observed to vary slightly between experiments (Figure S4I). We found that this model (Figure 4D) and its underlying assumptions can accurately describe both the effects of varying TF concentration and number of binding sites on the gene expression dynamics of the ARGFX bifunctional domain (Figure 4A-B, solid lines). As expected from our experimental observations, the model predicts that the rate of activation for the ARGFX bifunctional domain is fast and saturates at relatively low dox concentrations, while silencing occurs at a timescale two orders of magnitude slower than activation and requires high dox concentrations to contribute meaningfully to gene expression dynamics (Figure 4E).

To assess the ability of this integrated model to capture the behavior space of the ARGFX bifunctional domain, we used it to predict two summary metrics of our system for each dox concentration and number of binding sites: the difference between the maximum fraction of cells On and final fraction of cells On, a metric of repression strength (Figure 4F, G), and the final fraction of cells On, a metric of the steady state behavior (Figure 4H, I). At the minimal promoter, these parameters are together sufficient to describe the behavior of the gene expression response. The model can accurately describe the threshold at which repression begins to occur as a function of dox and number of TetO sites (Figure 4F, G). It also captures the non-monotonic, bell-shaped relationship between TF concentration (dox) or number of binding sites and the final fraction of active cells (Figure 4H, I). Notably, this non-monotonicity contrasts with canonical transcriptional activators such as VP48, which does not show occupancy-dependent repression (Figure S4E, J) and instead initially increases and then plateaus in its ability to activate as concentration and the number of binding sites increases (Figure 4H, I, pink; Figure S4J).

### Individual bifunctional domains can differentially regulate the expression dynamics of multiple genes in single cells using rationally designed regulatory elements

Together, our experiments and modeling revealed that the relative strengths of activation and repression by bifunctional domains is determined by domain occupancy at regulatory regions, which is set by a number of both *cis* (such as binding site number) and *trans* (such as TF concentration) variables. By independently tuning these parameters, individual cells could access a diverse set of dynamic gene expression responses using a single bifunctional domain. To better understand the behavior space of bifunctional gene expression dynamics at a minimal promoter in terms of these variables, we used our integrated model to predict gene expression dynamics for many combinations of binding site numbers and effective TF concentrations. From the predicted gene expression dynamics, we extracted the maximum fraction of active cells and the difference between this maximum and the steady state fraction of active cells to quantify the response (Figure 5A). From this analysis, we can define two functional regimes for bifunctional domains at the minimal promoter: an activation regime at combinations of binding site number and TF concentration that result in low DNA occupancy, and a pulse-generating regime where these variables combine to produce high DNA occupancy. In principle, multiple target genes in a single cell could lie in separate behavioral regimes by having different *cis* regulatory features, and therefore be regulated by the same bifunctional TF with distinct gene regulatory dynamics (Figure 5A, yellow vs. green dots in top box and associated curves on the top right). In addition, by changing a global variable such as TF concentration, these target genes could move between regions of the behavior space and be regulated with a new set of gene regulatory dynamics with the same domain (Figure 5A, yellow vs. green dots in lower box and associated curves on the bottom right).

**Figure 5:**
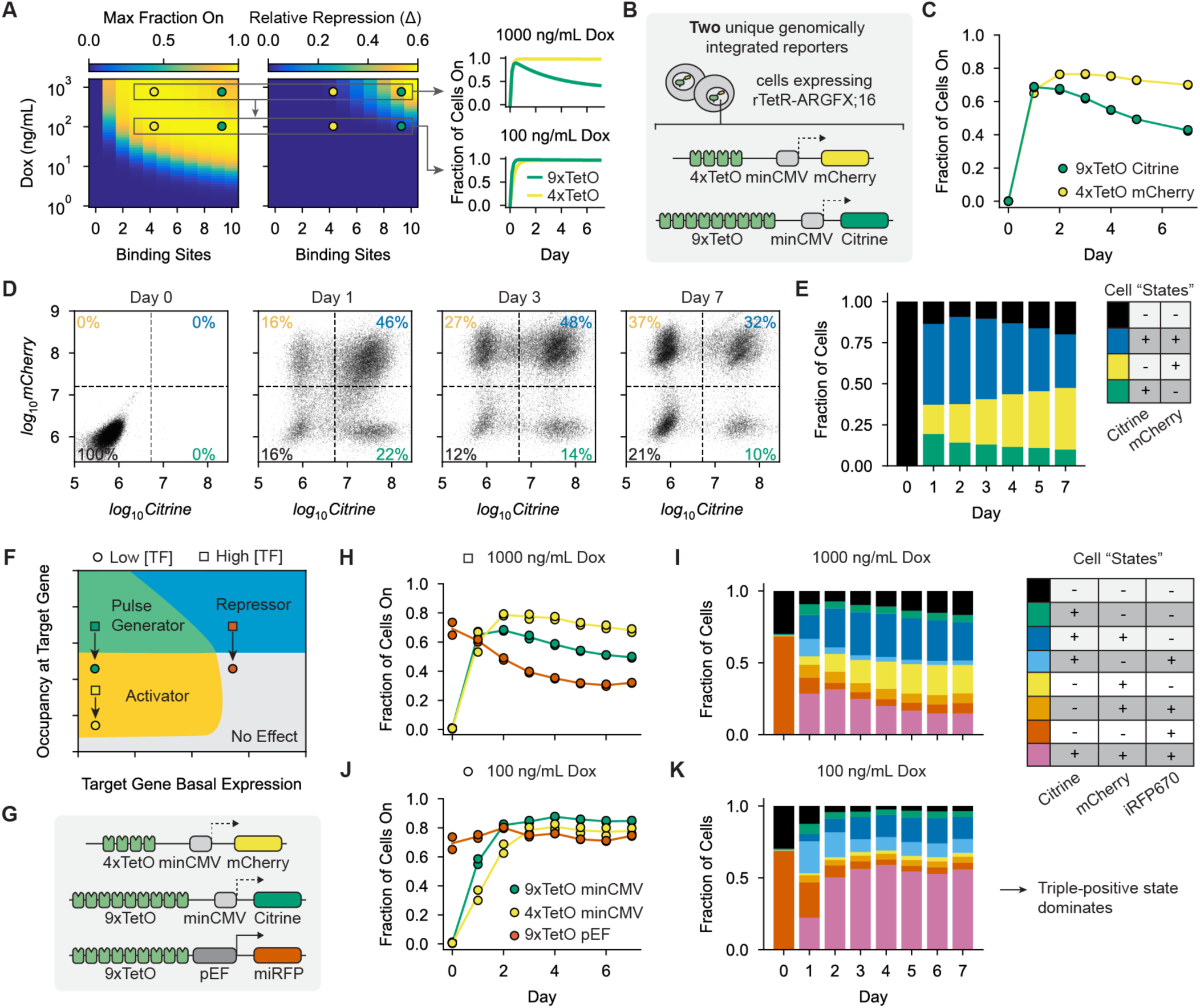
Rationally designed regulatory elements enable differential control of multiple genes in individual cells using a single bifunctional domain. **A)** Color intensities show model predictions (using parameters from Figure 4) for (left) the maximum fraction of active cells and (middle) the difference between this maximum and the final fraction of active cells (relative repression) for ARGFX;16 recruited to minCMV at different dox levels and number of TetO binding sites. (Right) Example predicted gene expression dynamics are shown for two genes with high (9x) and low (4x) numbers of binding sites in cells with high (1000ng/mL) and low (100 ng/mL) dox. **B)** Schematic of the reporter genes integrated into separate copies of the AAVS1 locus in K562 cells to create a dual-reporter cell line. **C)** Fraction of cells expressing each reporter during ARGFX;16 recruitment in the dual-reporter cell line. **D)** Scatterplot of mCherry vs. Citrine fluorescence for individual cells on Days 0, 1, 3, and 7 of ARGFX;16 recruitment. Dotted lines indicate gates in each channel, and the percentage of cells in each quadrant is noted in the corners. **E)** Dynamics of the fraction of cells in each reporter expression state over the duration of ARGFX;16 recruitment. **F)** Complete behavior space of bifunctional domains generated using the procedure in Figure S5E. The approximate locations of the three reporters in G-K are shown at high (squares) and low (circles) dox. **G)** Schematic of the reporter genes integrated into separate copies of the AAVS1 locus in K562 cells in the triple-reporter cell line. **H, J)** Fraction of active cells for each reporter during recruitment with 1000 ng/mL (H) or 100 ng/mL (J) dox recruitment. **I, K)** Dynamics of the fraction of cells in each possible reporter expression state during ARGFX;16 recruitment with 1000 ng/mL (I) or 100 ng/mL (K) dox.

We wondered if we could engineer this predicted multimodal regulation using a single bifunctional effector domain. We integrated two reporter genes into separate copies of the AAVS1 safe-harbor locus in human K562 cells expressing rTetR-ARGFX;16. Based on our model, we designed one reporter to contain 9 TetO sites upstream of the minCMV promoter driving Citrine expression, which should generate a pulse of gene expression at high dox, and the other to contain 4 TetO sites upstream of minCMV driving mCherry expression, which should stay active over the recruitment duration (Figure 5A, top right, Figure 5B). We verified that changing the reporter color did not affect the behavior of the bifunctional domains in single-reporter lines (Figure S5A). We then recruited ARGFX;16 in this dual-reporter cell line with high dox (1000ng/mL) for seven days. As predicted, the 4xTetO-mCherry reporter remained active on the population level over the duration of recruitment while the 9xTetO-Citrine reporter first activated and then silenced in the cell population (Figure 5C). At the single cell level, we observed all four possible expression combinations of the two reporters and that their relative abundances changed over time as expected for independent regulation of each reporter gene by ARGFX;16 (Figure 5D; Figure S5B). The four possible gene expression combinations in this dual-reporter cell line can be thought of conceptually as four “cell states.” We observed that a single bifunctional domain is sufficient to create and dynamically regulate the relative abundance of these four cell states with distinct kinetics - in this case, the fraction of double-positive cells decreases in the population over time as the fraction of mCherry-only cells increases as this reporter is stably activated while Citrine is silenced (Figure 5E).

In general, a bifunctional domain regulating *n* target genes can generate 2^#^gene expression states (+/-expression of each target gene) and dynamically regulate their fractional abundances in the cell population. Critically, the number of possible gene expression states does not depend on the basal expression level of target genes, as bifunctional domains can both activate and repress. This contrasts with activation and repression domains which can only regulate target genes unidirectionally and with monotonic kinetics (Figure S5C-D), i.e. a repressor binding at a target gene with no basal expression would have no effect. Therefore, bifunctional domains have access to a much richer behavior space than classic activation and repression domains. By combining different levels of DNA occupancy at target genes with different basal expression levels of these target genes, individual bifunctional domains can act as pure activators, pure repressors, or pulse-generators in single cells (Figure 5F; Figure S5E).

To explore this theoretical behavior space, we generated a new cell line with 3 reporter genes predicted to lie in the different regions of the ARGFX;16 behavior space: activating (4xTetO-minCMV-mCherry), repressing (9xTetO-pEF-miRFP670nano3), and pulse-generating (9xTetO-minCMV-Citrine) at maximum occupancy (Figure 5F-G). We recruited ARGFX;16 with high dox (1000ng/mL) and measured Citrine, mCherry, and miRFP670nano3 expression over 7 days. As expected, over the course of ARGFX;16 recruitment we observe silencing at 9xTetO-pEF reporter, a pulse at 9xTetO-minCMV reporter, and mainly activation at 4xTetO-minCMV reporter (Figure 5H; Figure S5F). A single bifunctional domain can therefore control the temporal dynamics of 8 “cell states,” represented by unique combinations of the three fluorescent reporters (Figure 5I). As expected, states with miRFP670nano3 active decrease in relative abundance over time, states with mCherry active increase, and states with Citrine active first increase then decrease (Figure 5I).

To this point, we have controlled domain occupancy at the three reporter genes only by changing *cis* variables (binding site number). Changing a *trans* parameter, such as domain concentration, would affect the occupancy of the bifunctional domain at all three reporters simultaneously. In some cases, this could result in a change in the behavior of the bifunctional domain at one or more reporters and result in a different set of kinetic responses. For example, reducing domain concentration in cells containing the three reporters could shift ARGFX;16 to acting as a pure activator at the 9xTetO-minCMV reporter and having no effect at the 9xTetO-pEF reporter (Figure 5F, circles). To demonstrate this principle, we recruited ARGFX;16 in our triple-reporter cell line with a lower dox concentration (100 ng/mL). As predicted, we observed strong expression from all three reporters over the duration of ARGFX;16 recruitment (Figure 5J). While the same 8 cell states were observed as with high dox, the relative composition of states over time changed drastically: at low dox, the state with all three reporters active comprises the majority of the population and grows over time (Figure 5K). Together, these data show that bifunctional domains allow for an extraordinary degree of economy and flexibility in gene regulation, which can be co-opted in engineered gene expression systems to create and dynamically regulate multiple gene expression states with a single input.

## Discussion

How cells regulate transient and adaptive transcriptional responses to signals is a central question of systems biology. Typically, feedback loops in gene regulatory networks or signaling pathways regulate pulses of gene expression(*57*, *58*). However, in this work we have shown that pulses of gene expression can also be intrinsically produced during constant recruitment of a specific class of bifunctional transcriptional effector domains without the need for more complex regulatory networks. Bifunctional effector domains are a recently discovered class of human transcriptional ED defined by their ability to activate a minimal promoter and repress a constitutively active promoter in an otherwise identical genetic context, and are found in dozens of human TFs and chromatin regulators(*21*). Using a high-throughput method to measure the expression dynamics of >1000 EDs, we found that pulse generation is a general feature of bifunctional effector domains (Figure 1, Figure S1). In addition, we observed that this ability is unique to bifunctional domains and is not shared by most classical activation domains, suggesting that pulse generation is enabled by the ability of bifunctional domains to also actively repress transcription, which is not shared by canonical activation domains. The gene expression dynamics enabled by bifunctional domains represent a new logic of transcriptional control accessible to human TFs and provide powerful tools for creating synthetic gene circuits with complex expression dynamics from a minimal set of components.

Using degraders or chemical inhibitors for various transcriptional cofactors, we observed that both coactivator and corepressor function is necessary for bifunctional domains to generate pulses of gene expression and that these domains induce distinct, temporally separated chromatin states defined by active and repressive chromatin modifications (Figure 2, Figure S2). These results are consistent with a minimal model where bifunctional domains recruit both coactivators and corepressors, either directly or indirectly, to affect activation and repression respectively. In such a model, a separation of timescales between fast coactivator and slow, dominant corepressor function results in a pulse of gene expression.

While it is known that certain TFs, such as Sp3, differentially interact with coactivators and corepressors through separate ADs and RDs depending on signaling or chromatin context (*59*, *60*), our results show that individual bifunctional domains can directly or indirectly recruit both types of cofactors to produce pulses of gene expression in a fixed context. This observation is similar to previous observations for the transcriptional activator White Collar Complex in *Neurospora*, which has been shown to rapidly recruit coactivators and induce histone acetylation and gene activation, then subsequently recruit histone deacetylases to repress transcription while remaining bound(*42*). Interestingly, while the basic mathematical form of pulse-generation associated with bifunctional regulation seems general, the specific cofactors necessary for repression varied widely across bifunctional domains (Figure 2G). Much work remains to be done in mapping domain-cofactor interactions for bifunctional domains found in diverse TF families.

Our synthetic expression system also allowed us to carefully dissect the dose dependence of pulse-generating bifunctional domains on their resulting gene expression dynamics. We found that pulse dynamics are controlled by the occupancy of bifunctional domains at the regulatory region of the reporter gene (Figure 4, Figure S4). At low occupancy bifunctional domains act only as activators. Past a certain occupancy threshold, the rate of repression gradually becomes stronger, producing pulses of expression and a non-monotonic relationship between TF input and steady state gene expression output. An integrated thermodynamic and kinetic model of gene expression can predict the effects of changing TF concentration and the number of TF binding sites on gene expression dynamics. Fits of this model to our experimental data revealed that the coactivators and corepressors recruited by bifunctional domains have different sensitivities to TF occupancy, with repression requiring a higher occupancy. This observation suggests that repression is the result of cooperative recruitment of corepressors, bifunctional domains have a lower affinity to corepressors than to coactivators, or that corepressors must be recruited in higher numbers to meaningfully affect gene expression dynamics. The specific biochemical mechanism causing the difference in coactivator and corepressor sensitivity to domain occupancy requires further investigation.

Other recent works have found examples of TFs that have a similar non-monotonic relationship between steady state gene expression and the number or affinity of TF binding sites upstream of a promoter(*44*, *61–63*). While these works did not measure gene expression dynamics to test if these TFs can produce the type of pulses described here, the steady state behavior they observed is consistent with our findings. Our experimental and theoretical work suggests that the non-monotonic behavior is a general principle of bifunctional TFs that either containing bifunctional effector domains (overlapping AD and RD, as is the case with our synthetic rTetR-EDs) or TFs with separate ADs and RDs. Previous studies have also proposed various models of gene expression that can predict this non-monotonicity(*43*, *64*). In general, these models posit that non-monotonicity occurs if a TF has opposing effects on two different steps of transcription (for example, it promotes RNA polymerase binding but inhibits elongation). However, the connections between these models and molecular mechanisms of transcriptional regulation are unclear and remain to be tested. In addition, these previous works do not consider or measure the gene expression dynamics produced by these bifunctional TFs. The single-cell resolution of our measurements over time reveals that bifunctional domains catalyze transitions between long-lived discrete gene expression states (On or Off that can be detected at the protein level) by dynamically modifying the chromatin state of target genes. This is contrary to predictions of existing models where repression acts on short-lived transcriptional steps that would predict a shift in the entire gene expression distribution. This suggests that models of pulse-generation and bifunctionality need to explicitly integrate chromatin regulation.

The potential for pulse-generating bifunctional domains to be used in engineering synthetic gene circuits with complex expression dynamics is particularly exciting. Current methods for engineering complex expression dynamics such as pulses require multiple TFs co-regulating each other and target genes to be integrated into individual cells. This approach suffers from limitations in efficient transgene integration in desirable loci(*65*), background silencing of transgenes(*32*) (which can be exacerbated the more genes that are integrated into individual cells), and cellular fitness defects arising from the overexpression of multiple proteins(*66*). By intrinsically regulating pulses of gene expression, bifunctional domains can produce desirable gene expression dynamics with only a single TF input, drastically simplifying this engineering problem.

In addition, because bifunctional domains function at the level of chromatin and their behavior is encoded by DNA occupancy, a single bifunctional domain can be used to regulate multiple target genes in individual cells with rationally engineered dynamics (Figure 5, Figure S5). Here, we have demonstrated that a single bifunctional domain can simultaneously activate one gene, repress a second, and produce a pulse of gene expression at a third in individual cells. Functionally, the domain produces 8 discrete “cell states” with a single input and dynamically regulates their relative abundance in the cell population over time in a predictable way. These bifunctional domains could be used in applications such as directed differentiation protocols, where certain genes must be activated, repressed, and transiently expressed over a timescale of days, though more work is required to test and validate their performance in various systems.

Overall, in this work we have dissected the mechanisms of pulse generation by bifunctional transcriptional effector domains and developed a quantitative framework for predicting and engineering this type of transcriptional control. Effector domains that intrinsically regulate pulses of gene expression represent a new logic of transcriptional control that must be taken into account when analyzing how TFs containing these domains fit into endogenous gene regulatory networks(*67*). These domains could also provide powerful tools for engineering compact synthetic circuits that display complex gene expression dynamics, a major goal in synthetic biology(*32*, *33*, *65*, *66*). The mechanistic and quantitative framework for understanding the gene regulatory dynamics of pulse-generating bifunctional domains developed here should serve as a foundation for these pursuits.

## Acknowledgments

We thank Alistair Boettiger for helpful discussions and constructive advice; Sage Allen for keeping the laboratory stocked and running; Abby Thurm for assistance with HT-Recruit and CUT&RUN protocols and data analysis as well as feedback on the manuscript; Saman Tabatabaee for feedback on the manuscript; and all members of the Bintu lab for advice and feedback. This work was supported by the Toyota Riken Overseas Scholarship (MS), JSPS Overseas Research Fellowship (TF), ARCS Foundation Fellowship (NVD), NIGMS T32GM141828-01A1 (CA), NIH T32 GM007276 (EJC), NIH 5T32HG000044-28 (GLJ), NSF GRFP DGE-1656518 (NVD), NIH MIRA R35GM128947 (LB), and NIH 4DN U01DK127419 (LB).

## Author contributions

LB, CJA, EJC, and NVD conceived the study. CJA and EJC performed experiments and analyzed data with assistance from GLJC, NVD, and MS. CJA and EJC performed mathematical modeling, with input from LB and TF. CJA, EJC, and LB wrote manuscript and prepared figures with feedback from all authors. LB supervised the study and acquired funding.

## Data and materials availability

Raw data related to HT-Recruit and CUT&RUN assays will be deposited on GEO. All reporter plasmids and recruiter constructs will be made available at Addgene or upon request. All data is available in the main text or the supplementary materials.

## Supplementary Materials

Materials and Methods Supplementary Text Figures S1 to S5

Tables S1 to S4 References (*68–76*)

