## Supplementary Information for "Bifunctional transcriptional effector domains control gene expression pulses in an occupancy-dependent manner"

**The PDF file includes:**

Supplementary Text
Materials and Methods
Figures S1 to S5
References (68-76)

### Supplementary Text

### Mathematical Model of Bifunctional Gene Regulation

*Three states of gene expression during bifunctional domain recruitment*

Because we observed discrete, bimodal gene expression states during bifunctional domain recruitment at the minCMV, PGK, and pEF promoters (Figure 1B-D; Figure S4A), and bifunctional regulation relies on the recruitment of cofactors that modify chromatin state (Figure 2), we adopted a phenomenological, state-based modeling approach previously used to describe gene expression dynamics following the recruitment of epigenetic repressors(*68*, *69*). In this approach, cells are assumed to exist in a number of discrete gene states that include both promoter activity and its chromatin state (active, silenced, or ground state where the promoter is not transcribed but not repressed by silenced chromatin modifications) and transition rates between states are determined by TFs and chromatin remodelers.

This framework is not commonly used to model transcriptional activation by TFs, and bimodal expression levels are typically attributed to bursty transcription where the timescale of switching between promoter states is longer than the half life of mRNA and protein(*5*). However, our sorting experiments (Figure 3) reveal that the active and silenced gene expression states created by bifunctional domains are very long lived, otherwise re-equilibration would occur between these cell populations. Our state-based approach is essentially a limit of a promoter-switching model where transitions are very infrequent, and the level of TF binding controls the fraction of cells that transition into the active state. The level of TF binding once cells are in the active state likely also affects the average rate of transcription from the reporter gene, as predicted by thermodynamic models of gene regulation by transcriptional activators(*70*, *71*). Similar to the fraction of active cells, we see a non-monotonic relationship between dox concentration and number of binding sites and the mean Citrine intensity (MFI) of the active cells (Figure S4A). Here, as we develop and fit the theoretical model, we primarily consider the fraction of cells in the active state rather than the MFI of the active cells as this is the major change we see in the cell population over time.

Because three discrete gene expression states were directly observed at the PGK promoter (Figure S3A) and because the Citrine-Off cells after ARGFX;16 recruitment are enriched for repressive histone modifications compared to Citrine-Off cells before ARGFX;16 recruitment (Figure 2E), we initially adopted a model with three discrete gene expression states: a ground state (G), whose level of Citrine expression is determined by the promoter, an active state (A), where Citrine expression is increased over the baseline level, and a silent state (S) where the locus is actively repressed and no Citrine expression occurs. Because there is no epigenetic memory of silencing after dox washout and releasing ARGFX;16 from the reporter (Figure 1E), we include only one silent state. The ground state varies in gene expression and histone modifications between promoters: for example, for minCMV the ground state has an inactive promoter but lacks any active or repressive histone modifications (Figure 2D, E), while for pEF the ground state is transcriptionally active and has high levels of active histone modifications(*72*, *73*)

Based on our chemical inhibitor data (Figure 2), we assume that bifunctional domains interacting with coactivators influence transitions to the active state and domains interacting with corepressors influence transitions to the silent state.

*A three-state kinetic model of bifunctional gene regulation*

To start modeling the gene regulatory dynamics of bifunctional domains, we constructed a simple kinetic model where cells start in the ground state and can transition to both the active state and the silent state (Figure 1J; Figure 2A; Figure S1M-N). We assume that transitions are irreversible, that all cells in the active state can be silenced (transition to the silent state), and that the transition rates from ground to silent and from active to silent are equal. This model can be represented as a system of three differential equations:

$$\frac{dG}{dt} = -(k_1 + k_2)G(t) \tag{1}$$

$$\frac{dA}{dt} = k_1 G(t) - k_2 A(t) \tag{2}$$

$$\frac{dS}{dt} = k_2 (G(t) + A(t)) \tag{3}$$

Where *G(t)* is the fraction of cells in the ground state, *A(t)* the fraction of cells in the active state, *S(t)* the fraction of cells in the silent state, $k_1$ is the activation rate, and $k_2$ is the rate of silencing. Assuming that all cells begin in the ground state, these equations have the following analytic solution:

$$G(t) = e^{-(k_1+k_2)t} \tag{4}$$

$$A(t) = e^{-k_2 t} - e^{-(k_1+k_2)t} \tag{5}$$

$$S(t) = 1 - e^{-k_2 t} \tag{6}$$

From these equations (Equations 4-6), it is clear that at steady state ($t \rightarrow \infty$), $A = 0, G = 0$, and $S = 1$, meaning that all the cells are predicted to eventually be silenced.

We can also determine the amplitude of the pulse and the timing of its peak from the analytical solutions (Equations 4-6). We find the time of the maximum fraction of cells in the active state by taking the first derivative of *A(t)* with respect to time:

$$A'(t) = -k_2 e^{-k_2 t} + (k_1 + k_2)\, e^{-(k_1+k_2)t} \tag{7}$$

By setting the derivative equal to 0, we find that *A(t)* has either a maximum or minimum point at:

$$t^* = t_{A\prime=0} = \frac{1}{k_1} ln\left(\frac{k_1 + k_2}{k_2}\right) \tag{8}$$

To determine if *A(t)* is at a maximum or minimum at this time, we can take the second derivative of *A(t)*:

$$A''(t) = k_2^2 e^{-k_2 t} - (k_1 + k_2)^2\, e^{-(k_1+k_2)t} \tag{9}$$

Evaluating the second derivative at $t = t^*$, we obtain:

$$A''(t^*) = -k_1 k_2 \left(\frac{k_1 + k_2}{k_2}\right)^{\frac{-k_2}{k_1}} \tag{10}$$

If $k_1 > 0$ and $k_2 > 0$ (which must be true for physical solutions), the second derivative is negative at this point, indicating that *A(t)* always has a maximum value before approaching 0 and therefore always produces a pulse of gene expression. However, only some of these pulses will occur over the timescales of our experiments and with large enough amplitudes to detect experimentally. For this to occur, the rate of activation ($k_1$) must be sufficiently large to activate the majority of cells and the rate of repression ($k_2$) must be in a "sweet spot" where the cells will be silenced after a few days of recruitment, but not so large that it prevents cells from activating (Figure 1L; Figure S1M-N).

*Models that are consistent with a non-zero steady state fraction of active cells*

We have observed that a small but stable fraction of cells remained in the active state even when recruiting ARGFX;16 to the minCMV promoter for over two weeks (Figure 3A), which is inconsistent with the minimal model of bifunctional pulse generation described above that predicts all cells silence at steady state. Therefore, we needed to update our simple, three-state model to better understand this phenomenon. We reasoned that two minor extensions of this three-state model could account for the non-zero steady state fraction of active cells: a model capable of reaching a dynamic equilibrium between the states, and a model with an additional terminal state of active cells that cannot be silenced (Figure 3B).

In order for the system to reach a dynamic equilibrium, a back rate from the silent state to either the active or ground states must be present. In the most general case, there are six possible transitions between the three gene expression states in our model (Figure S3B). This model can be represented by the following system of differential equations:

$$\frac{dG}{dt} = k_3A(t) + k_5S(t) - (k_1 + k_2)G(t) \tag{11}$$

$$\frac{dA}{dt} = k_1G(t) + k_4S(t) - (k_3 + k_2)A(t) \tag{12}$$

$$\frac{dS}{dt} = k_2(G(t) + A(t)) - (k_4 + k_5)S(t) \tag{13}$$

To determine the steady state behavior of the system, we use the constraint $A + G + S = 1$, set the differential equations equal to 0, and solve for the fraction of cells in each state. We obtain:

$$G_{ss} = \frac{k_3k_4 + k_2k_5 + k_3k_5}{(k_1 + k_2 + k_3)(k_2 + k_4 + k_5)} \tag{14}$$

$$A_{ss} = \frac{k_1k_4 + k_2k_4 + k_1k_5}{(k_1 + k_2 + k_3)(k_2 + k_4 + k_5)} \tag{15}$$

$$S_{ss} = \frac{k_2}{k_2 + k_4 + k_5} \tag{16}$$

Because each rate parameter must be greater than 0, we see that each state accounts for a non-zero fraction of the cell population in steady state. This fraction of cells does not depend on the initial condition of the system (i.e. a population of cells starting 100% in the active state will reach the same steady state as a population of cells starting 100% in the ground state, Figure S3C). To illustrate this phenomenon, we can fit this model to the measured fraction of cells expressing Citrine over time for 10 days where all cells start in the ground state (Figure 3A, Figure S3B), and then use the fit parameters to predict gene expression dynamics of cell populations starting in each of the other two cell states (all active or all silent, Figure S3C). As expected, the final steady state of active cells is independent of the starting population and cells are predicted to dynamically re-equilibrate to these steady state values after sorting. However, we do not observe dynamic re-equilibration when we sort the cells and propagate them (Figure 3C).

An alternative model is that a fourth gene expression state exists where cells express Citrine but are unable to transition to the silent state, which we refer to as the sustained active state ($A*$) (Figure 3B, right; Figure S3D). Intuitively, this system has both active and silent "attractor" states that cells cannot escape from due to a lack of backwards transition rates. This system can be represented by the following system of differential equations:

$$\frac{dG}{dt} = -(k_1 + k_2)G(t) \tag{17}$$

$$\frac{dA}{dt} = k_1 G(t) - (k_2 + k_3)A(t) \tag{18}$$

$$\frac{dS}{dt} = k_2(G(t) + A(t)) \tag{19}$$

$$\frac{dA^*}{dt} = k_3 A(t) \tag{20}$$

We allow the transition rate to the sustained active state A*, $k_3$, to differ from the transition rate from the ground state to the active state, $k_1$, because we have no reason to think that the mechanism of initial activation is the same as the mechanism responsible for creating stably active cells. Even with the constraining equation$A + G + S + A^* = 1$, the steady state behavior of this system does not have a universal solution, as it depends on both the rate parameters and the initial conditions. For example, if 100% of cells started in the silent state, the steady state fraction of cells in the sustained active state would be 0. If 100% of cells started in the sustained active state, the steady state fraction would also be 100%. This is illustrated in Figure S3E.

For our experiments at the minCMV promoter, we can make the simplifying assumption that all cells start in the ground state ($G(0) = 1$). In this case, the system has the analytic solution:

$$G(t) = e^{-(k_1+k_2)t} \tag{21}$$

$$A(t) = \frac{-k_1}{k_1 - k_3}\left(e^{-(k_1+k_2)t} + e^{-(k_2+k_3)t}\right) \tag{22}$$

$$S(t) = \frac{k_2(k_1+k_2+k_3)}{(k_1+k_2)(k_2+k_3)} + \frac{k_2k_3}{(k_1+k_2)(k_1-k_3)}e^{-(k_1+k_2)t} - \frac{k_1k_2}{(k_1-k_3)(k_2+k_3)}e^{-(k_2+k_3)t} \tag{23}$$

$$A^*(t) = \frac{k_1k_3}{(k_1+k_2)(k_2+k_3)} + \frac{k_1k_3}{(k_1+k_2)(k_1-k_3)}e^{-(k_1+k_2)t} - \frac{k_1k_3}{(k_1-k_3)(k_2+k_3)}e^{-(k_2+k_3)t} \tag{24}$$

At steady state ($t \to \infty$), the fraction of cells in each state is:

$$G_{ss} = 0 \tag{25}$$

$$A_{ss} = 0 \tag{26}$$

$$S_{ss} = \frac{k_2(k_1+k_2+k_3)}{(k_1+k_2)(k_2+k_3)} \quad (27)$$

$$A^{*}_{ss} = \frac{k_1 k_3}{(k_1+k_2)(k_2+k_3)} \quad (28)$$

In Figure S3D-E, we demonstrate the dependence of the steady state behavior on initial conditions by fitting this model to the fraction of active cells during the first 10 days of ARGFX;16 recruitment (all cells starting in the ground state), then use the fit rate parameters to predict gene expression dynamics for populations of cells starting in each of the other cell states (sorted active, stably active or silent). As expected, populations starting in the sustained active and silent states remain in the states indefinitely during ARGFX;16 recruitment.

*Developing a thermodynamic binding model of bifunctional effector domains*

It is well established that the rates and levels of activation and repression depend on a number of parameters such as TF concentration and the number of TF binding sites at a target gene(*54–56*, *74*, *75*). Our dox titration and binding site perturbation experiments (Figure 4) suggest that these parameters also affect the dynamics of bifunctional gene regulation. In particular, at low dox concentrations and numbers of binding sites bifunctional domains act only as activators at the minimal promoter, while at high concentrations and numbers of binding sites they generate pulses of gene expression (Figure 4).

We wondered if we could extend our current model to take both the number of binding sites and TF occupancy as inputs and faithfully predict gene expression dynamics as output. To do this, we must use a framework that is able to connect microscopic variables such as binding site number and TF abundance with macroscopic changes in gene expression states. Thermodynamic models are powerful tools for predicting gene expression while accounting for these microscopic variables(*70*). These models assume that the system of interest can exist in a finite set of discrete microstates. If the system is in thermodynamic equilibrium, statistical mechanics can be used to calculate the probability of the system occupying any one microstate based on the relative energies of each state (which is determined by molecule concentrations, binding affinities, etc.) Because of their ability to directly account for multiple determinants of TF occupancy, we therefore developed a thermodynamic binding model of bifunctional domains.

Typically, thermodynamic models are used to predict the probability that RNA polymerase or other transcriptional machinery is bound at a promoter. It is assumed that there is a separation of timescales between TFs binding and unbinding and transcription, therefore the rate of transcription is proportional to the probability that RNA polymerase is bound and one can predict gene expression dynamics(*15*). Based on our observations, we hypothesize that TF binding and unbinding modifies the slow transitions between discrete gene expression or chromatin states in addition to influencing transcription rates directly. Because we are mainly interested in the overall gene state, we do not model RNA polymerase binding and focus only on the binding and unbinding of TFs and their effects on the gene states we have defined: ground, silent, active and stably active.

To develop a thermodynamic binding model of bifunctional domains, we assume that a single bifunctional domain can exist in two functional forms, depending on whether it is interacting with a coactivator ($TF_A$) or a corepressor ($TF_R$) (Figure 4D, left), and that the average occupancy of these different forms determines the transition rates to active and silent states respectively (Figure 4D, right). The concentrations of these two TF functional forms in part determine their average occupancy. To relate the relative abundance of each form to the dox concentration used to recruit the bifunctional domain to the reporter we use a simple first order binding reaction to model dox binding to rTetR:

$$[TF_A] = \frac{\theta[dox]}{K_{dox} + [dox]}) \tag{29}$$

$$[TF_R] = \frac{(1-\theta)[dox]}{K_{dox} + [dox]} \tag{30}$$

Where $K_{dox}$ is the dissociation constant of dox binding to rTetR and $\theta$ is the fraction of dox-bound TF that is interacting with the coactivator.

To determine average occupancy, it is useful to calculate the partition function of the system, which is obtained by summing the statistical weights of every possible microstate (Figure 4D). For a regulatory region with $N$ binding sites that can be independently bound by both active and repressive forms of the bifunctional domain, the partition function is:

$$Z = \sum_{n_A=0}^{N} \sum_{n_R=0}^{N-n_A} \frac{N}{n_A} \frac{N-n_A}{n_R} ([TF_A]K_{TF})^{n_A}([TF_R]K_{TF})^{n_R} = (1 + K_{TF}[TF_A] + K_{TF}[TF_R])^N \tag{31}$$

Where $K_{TF}$ is the association constant of the transcription factor, which we assume to be the same for the activating and repressing forms (since they are both fused to rTetR), and $n_A$ and $n_R$ is the number of activator and repressor forms bound, respectively. We assume there is no binding cooperativity between the domains for simplicity. We can then calculate the probability of a certain number of molecules being bound by summing the statistical weights of states with that number of molecules bound and dividing by the partition function:

$$P(n_A) = \frac{1}{Z} \sum_{n_R=0}^{N-n_A} \frac{N}{n_A} \frac{N-n_A}{n_R} ([TF_A]K_{TF})^{n_A}([TF_R]K_{TF})^{n_R} = \frac{1}{Z} \frac{N}{n_A} ([TF_A]K_{TF})^{n_A}(1 + [TF_R]K_{TF})^{N-n_R} \tag{32}$$

$$P(n_R) = \frac{1}{Z} \sum_{n_A=0}^{N-n_R} \frac{N}{n_R} \frac{N-n_R}{n_A} ([TF_A]K_{TF})^{n_A}([TF_R]K_{TF})^{n_R} = \frac{1}{Z} \frac{N}{n_R} ([TF_R]K_{TF})^{n_R}(1 + [TF_A]K_{TF})^{N-n_A} \tag{33}$$

Then, the average number of each molecule bound at the regulatory region is given by:

$$\langle TF_A \rangle = \sum_{n_A = 0}^{N} P(n_A) n_A = N \left( \frac{K_{TF}[TF_A]}{1 + K_{TF}[TF_A] + K_{TF}[TF_R]} \right) \quad (34)$$

$$\langle TF_R \rangle = \sum_{n_R = 0}^{N} P(n_R) n_R = N \left( \frac{K_{TF}[TF_R]}{1 + K_{TF}[TF_A] + K_{TF}[TF_R]} \right) \quad (35)$$

In summary, we have obtained expressions for the average occupancy of bifunctional domains interacting with coactivators vs corepressors as a function of the number of binding sites in the reporter gene and the dox concentration used to recruit the domains to the reporter.

*Integrating a thermodynamic binding model with a kinetic model of bifunctional regulation*
A central assumption of our model is that the average occupancy of the activator and repressor forms of the bifunctional domain determines the transition rates between discrete gene expression states in our kinetic model. Because we do not know *a priori* what the functional form of the relationships between occupancy and the transition rates is, we model each transition rate as a Hill function dependent on average occupancy (Figure S4G). For the divergent states model that is consistent with our sorting data (Figure S4H, this becomes:

$$k_{1,3} = \frac{k_{1,3\,max} \ \langle TF_A \rangle^{h_{1,3}}}{{D_{1,3}}^{h_{1,3}} + \langle TF_A \rangle^{h_{1,3}}} \quad (36)$$

$$k_2 = \frac{k_{2\,max} \ \langle TF_R \rangle^{h_2}}{{D_2}^{h_2} + \langle TF_R \rangle^{h_2}} \quad (37)$$

Where $k_{max}$ is the maximum value of a transition rate, $h$ is the Hill exponent, and $D$ is the TF occupancy at which the transition rate is half maximal. Because we noticed slight variations in the strength of activation and repression between our experiments, which were conducted using cell lines of different ages and that had been freeze/thawed different numbers of times, we allowed the $k_{max}$ parameters to be fit individually for each dataset. All other parameters were fit globally.

We initially tried to fit our data to the simple, Divergent States model shown in Figure 3B and S4H. This model performed well at predicting gene expression dynamics for intermediate and high numbers of binding sites and dox concentrations, but failed to predict dynamics at very low numbers of binding sites and dox concentrations where no repression occurs but activation is so weak that only a fraction of cells activate (Figure S4H, yellow). We hypothesize that this sub-maximal fraction of cells that activate is caused by cells remaining in the ground state at lower TF occupancy, which is not accounted for in our model and assumes that all cells will eventually transition out of the ground state.

To account for sub-maximal activation, we included a back rate from the active to ground state that was assumed to be occupancy-independent (Figure 4D, right; Figure S4I). This updated model was able to successfully fit both our binding site and dox perturbation datasets (Figure 4A-B; Figure S4I) and could describe well the behavior space of the bifunctional domains, predicting the occupancy threshold at which the rate of repression begins to increase and the non-monotonic relationship between occupancy and steady state gene expression levels (Figure 4E-I).

To compare this predicted behavior space with that of the classic transcriptional activator VP48, we fit VP48 dox titration and binding site perturbation to a similar, but modified, model (Figure S4E, J). Because VP48 does not actively repress genes, any slight decreases in gene expression over time are likely caused by background silencing. Therefore, we assumed that the transition rate to the silent state was occupancy independent. This model could successfully describe the VP48 data, and showed a monotonic relationship between the fraction of active cells and domain occupancy (Figure 4H and I, pink), as expected for classical activation domains.

It is important to note that we are accounting for nonlinearity between TF abundance and transition rates between gene states through these Hill functions and do not model any sort of binding cooperativity between TFs, despite this type of cooperativity being observed for a handful of activation domains(*56*). It is likely that a similar fit could be achieved by modeling cooperative binding between TFs and assuming a linear relationship between TF occupancy and transition rates between gene expression states.

**Materials & Methods**

**Plasmid cloning**

Effector domains were ordered as gBlocks from IDT and cloned as a fusion with rTetR using GoldenGate cloning in the backbone pJT126 (Addgene no. 161926), backbone CA024 (Figure S1D), or backbone EC051 (Figures 5, S5). Golden Gate assembly was performed in 10uL reactions using 80ng of plasmid backbone, 5ng of insert (2:1 molar ratio of insert/backbone), 1uL of 10x T4 DNA Ligase Buffer (NEB, B0202S), 0.25uL T4 DNA Ligase 2,000,000U/mL (NEB, M0202T), 0.75uL Esp3I restriction enzyme (NEB, R0734L), and nuclease-free ddH2O. Typically, 35 rounds of digestion-assembly cycles were performed in a thermocycler (Bio-Rad).

To construct backbone CA024, the 3xFLAG sequence between rTetR and the effector domain was replaced with HaloTag using Gibson assembly. To construct backbone EC051, the T2A-mCherry-BSD sequence in plasmid CA024 was replaced with a T2A-BSD gene fragment ordered from IDT also using Gibson assembly. The minCMV reporters with different numbers of binding sites (plasmids EC040-EC048; Figures 4, S4) were designed by sequentially scrambling the sequences of TetO sites sequentially from the 5' end of the 9xTetO minCMV reporter plasmid (Addgene no. 161928). These new binding site arrays were ordered in split halves in the middle of the binding region as gene fragments from TWIST and assembled into backbone CL056 using three-part Gibson assembly. The reporters used for the multi-output experiments (Figures 5 and S5) were engineered by replacing the Citrine in the pEF reporter (Addgene no. 161927) and EC044 with either mCherry or miRFP670nano3 using Gibson assembly.

For all Gibson assemblies, vectors were obtained using restriction enzyme digestion of the relevant backbone followed by gel extraction (Zymo, D4001T). Insert fragments were obtained using PCR from plasmid templates followed by gel purification or ordered as gene fragments from IDT or TWIST. 20-30bp of overhangs were used for all assemblies. Reactions were performed in 20uL total with 50-100ng of gel extracted vector, a 2-fold molar excess of insert DNA (resuspended gene fragments or PCR product), and 10uL of NEBuilder HiFi DNA Assembly Master Mix (NEB, E26211L). Reactions were incubated at 50C for 60 minutes, then 2uL of assembly product was transformed into Mix and Go chemically competent cells. All cloned constructs were verified using whole-plasmid sequencing (Plasmidsaurus).

**K562 cell culture and cell line engineering**

All experiments presented here were performed in K562 cells (ATCC, CCL-243, female) except for those in Figure S1C-D. Cells were cultured in a controlled humidified incubator at 37C and 5% $CO_2$, in RPMI 1640 (Gibco, 11-875-119) media supplemented with 10% fetal bovine serum (FBS) (Takara, 632180) and 1% penicillin streptomycin (Gibco, 15-140-122). Cells were kept at a confluency of less than 1 million cells/mL of culture. HEK293T-LentiX (Takara Bio, 632180, female) cells, used to produce lentivirus (described below), were grown in DMEM (Gibco, 10569069) media supplemented with 10% FBS (Takara, 632180) and 1% Penicillin Streptomycin Glutamine (Gibco, 10378016).

minCMV, PGK, and pEF reporter cell line generation was performed as previously described(*21*, *35*). Briefly, pEF (Addgene no. 161927), minCMV (Addgene no. 161928), and PGK (Addgene no. 196545) promoter reporter cell lines were created by TALEN-mediated homology-directed repair to integrate donor constructs into the *AAVS1* locus by electroporating 1 million K562 cells with 1000 ng of reporter donor plasmid and 500 ng of each TALEN-L (Addgene no. 35431) and TALEN-R (Addgene no. 35432) plasmid using program T-016 on the Nucleofector 2b (Lonza, AAB-1001). After 48 hours, the cells were treated with 500 ng/mL puromycin until all non-resistant cells were killed and a population where the donor was stably integrated in the intended locus remained (5-7 days). For the recruitment assay shown in Figure 1E, a clonal PGK reporter cell line was obtained by sorting single cells into individual wells of a 96-well plate using a Sony Cell Sorter SH800S.

Effector domain recruitment constructs were stably integrated into K562 cells using lentivirus as previously described(*21*). Briefly, HEK293T-LentiX (Takara #632180) were plated in 6-well tissue culture plates in 2mL of DMEM such that they would be at ~80% confluency the next day, grown overnight, and transfected with 750ng of an equimolar mixture of the three third generation packaging plasmids (pMD2.G: Addgene no. 12259; pRSV-Rev: Addgene no. 12253; pMDLg/pRRE: Addgene no. 12251, all gifts from Didier Trono) and 750ng of recruiter construct plasmid using polyethylenimine (Polysciences no. 23966). After 72 hours of incubation, lentivirus was harvested and filtered through a 0.45mm polyvinyl difluoride filter (Millipore). Reporter K562 cells were transduced with the lentivirus by spinfection for 2 hours using 1mL of harvested virus per 300k cells. After 48 hours, infected cells were selected with 10 μg/mL of blasticidin (Gibco) until at least 80% of cells expressed the recruiter construct, as measured by mCherry expression via flow cytometry (Bio-Rad ZE5).

To generate multi-reporter cell lines (Figures 5, S5), 5 million K562 cells were electroporated with 3ug of each reporter plasmid (JT039 and EC059 for the dual-pEF line, DY032 and EC057 for the 9x and 4x TetO minCMV line, and DY032, EC057, and EC060 for the triple-reporter line) and 500 ng of each TALEN-L (Addgene no. 35431) and TALEN-R (Addgene no. 35432) plasmid using program T-016 on the Nucleofector 2b (Lonza, AAB-1001). After 48 hours, cells were treated with 500ng/mL puromycin for 7 days. To construct the 9x and 4x TetO minCMV line, 10 million selected cells were transiently transfected with colorless rTetR-VP48 (EC052) and replated in 1000ng/mL dox. The following day, mCherry+ and Citrine+ cells were isolated using a Sony Cell Sorter SH800S. Colorless rTetR-VP48 (EC052), rTetR-ARGFX;16 (EC054), and ZNF10-KRAB (EC053) were stably integrated into the appropriate sorted cell lines using lentivirus. To construct the triple reporter line, selected cells were first sorted for miRFP670nano3+ cells and then spinfected with colorless rTetR-VP48 and rTetR-ARGFX;16 as described above. All cells were treated with blasticidin until the surviving cells doubled roughly every 24 hours in the presence of blasticidin, demonstrating resistance to the selection marker. To finish creating the triple reporter line, cells were treated with 100ng/mL dox for 24 hours and miRFP670nano3+, Citrine+, and mCherry+ cells were enriched by cell sorting.

All experiments were performed with at least two biological replicates at the level of spinfection or reporter line electroporation. These cell lines were not authenticated. All cell lines tested negative for mycoplasma.

**hiPSC cell culture and cell line engineering**

hiPSC WTc11 parent cell lines were thawed and seeded on GFR Matrigel (Fisher Scientific 356231)-coated plates in mTESR Plus media (STEMCell Technologies 100-0276) supplemented with ROCK inhibitor Y-27632 (STEMCell Technologies, 72308). Cells were cultured in a controlled humidified incubator at 37C and 5% CO2. After 24 hours, the media was changed to mTESR Plus. Media was replenished every 24 hours.

hiPSC cell lines containing the 9xTetO-minCMV reporter (DY032) were generated by transfecting 1 million cells with 1000 ng of reporter plasmid and 500ng of each TALEN using Lipofectamine LTX. After 48 hours, cells were treated with 500 ng/mL puromycin for 7 days. Lentivirus containing the rTetR-ARGFX;16 recruitment construct (NVD146) was generated as described above, then harvested and concentrated using Takara Lenti-X Concentrator (Takara 631231) according to manufacturer's instructions. minCMV reporter lines were reverse transfected and incubated for 48 hours before selection with 10 µg/mL blasticidin for 7 days.

**HEK293T cell culture and cell line engineering**

HEK293T-LentiX cells were cultured in DMEM supplemented with 10% FBS and 1% penicillin-streptomycin-glutamine, described above. Reporter cell lines containing the pEF reporter (JT039) were previously engineered through Lipofectamine LTX transfection of TALENs and reporter plasmid, then selected with 0.25 µg/mL puromycin. A clonal cell line was then generated(*76*). Lentivirus containing the ARGFX;16 recruitment construct (NVD146) was generated as described above, then harvested and reverse transfected in reporter cells. Cells were incubated for 48 hours before selection with 10 µg/mL blasticidin for 7 days.

**Individual recruitment assays and flow cytometry measurements**

To measure gene expression dynamics during effector domain recruitment, $4x10^5$ K562 reporter cells stably expressing the recruiter construct were typically plated in 1mL of media in separate wells of a 24-well plate and either treated with doxycycline (Fisher Scientific), chemical inhibitors, or untreated. Time points were measured by flow cytometry analysis (Bio-Rad ZE5, Everest v.2.3-3.0). Dox and inhibitors were assumed to degrade each day, so fresh dox/inhibitor media was added each day of the time course.

For recruitment assays with hiPSC reporter lines, cells were dissociated with Accutase and baseline Citrine expression was measured with flow cytometry. $4x10^4$ cells per well were seeded in 4 matrigel-coated 12-well replica plates in mTESR Plus supplemented with 10µM ROCK inhibitor and either 1000 ng/mL or 0 ng/mL dox. 2 replica wells were seeded per condition. During day 1 - 3 of dox recruitment, wells of one replica plate were dissociated with accutase and mCherry and Citrine fluorescence were measured by flow cytometry. Cells reached confluence on day 3, and so cells from the remaining replica plate were dissociated and re-seeded on fresh matrigel-coated plates for flow cytometry analysis on day 4 and 5 of dox recruitment. Media was replenished every 24 hours for the duration of the timecourse.

For recruitment assays with HEK293T reporter lines, cells were dissociated with Trypsin-EDTA (0.25%) and baseline $2.5x10^4$ cells per well were seeded in a 24-well plate in DMEM supplemented with 10% FBS and 1x Pennicilin-Streptomycin. 2 replica wells were seeded per dox condition (1000 ng/mL or 0 ng/mL) per day (5 days total). Cells were reverse plated such that each day, dox was added to additional wells so that each well had been cultured in dox media for 0 - 4 days. After 4 days of recruitment, cells were dissociated and flowed for Citrine expression.

For recruitment assays with cell sorting (Figures 3, S3; Sony Cell Sorter SH800S), K562 reporter cells stably expressing recruiter constructs were grown in larger volumes of media that were necessary to obtain sufficient cells after cell sorting. Throughout timecourses, cells were maintained at a confluency of 4-8x$10^5$ cells/mL and split 1:2 each day to remain consistent with our 24-well plate experiments. For sorting, cells were centrifuged at 300g for 5 minutes and the resuspended at a concentration of 10-20x$10^6$ cells/mL in doxycycline media. Collection tubes were prepared with doxycycline media to minimize the amount of time cells spent without dox during sorting. After a sufficient number of cells were sorted, collection tubes were spun down at 300g for 5 minutes and cells were replated at 400k cells/mL in doxycycline media.

**Flow cytometry analysis**

Data were analyzed using Cytoflow (v.1.1, https://github.com/bpteague/cytoflow) and custom Python scripts. Events were gated for viability and mCherry as a delivery marker for the recruiter constructs, except for the colorless recruiter constructs used in the multi-reporter experiments (Figures 5, S5). To compute the fraction of On cells during dox recruitment to minCMV, we fit a Gaussian model to untreated reporter-matched cells to model the distribution of Off cells, then set a threshold that was two standard deviations above the mean of the Off peak to label active cells. We do the same for computing the fraction of Off cells for the constitutively active pEF promoter, but fit a two component Gaussian mixture model to untreated reporter cells to describe background-silenced cells and set a threshold that is two standard deviations below the mean of the On peak. For PGK reporter cells, we fit a Gaussian model to untreated reporter cells and set thresholds both two standard deviations above the mean to call On cells and two standard deviations below the mean to call Off cells. For multi-reporter cell lines (Figures 5, S5), thresholds were set manually to be roughly halfway between the Off and On peak for each reporter construct, in log10 space. Thresholds are shown in Figure 5D and Figure S5G.

**Screen and validations for bifunctional domain dynamics**

An HT-recruit assay to measure gene expression dynamics following activation and bifunctional domain recruitment and subsequent library preparation was performed as described previously(*21*, *35*). We recruited the activating hits validation library from DelRosso *et al.*, 2023, containing 1,355 activating and bifunctional tiles as well as random tiles and negative, non-activating control tiles. The library was previously designed, cloned, and delivered to K562 cells containing the minCMV reporter gene as described in DelRosso *et al.,* 2023 (*21*). Samples were treated with 1000 ng/mL doxycycline for either 1 or 5 days and maintained at >1000x library coverage. Off and On populations were magnetically separated, genomic DNA was extracted, and library preparation was performed as previously described(*21*). Libraries were sequenced using an Illumina MiSeq with 2x150 cycles.

A logistic model, including a scale parameter, was fit to the validation and screen data using SciPy's curve fit function.

**Analysis of HT-Recruit data**

HT-Recruit data was processed and enrichment scores were computed using the HT-Recruit-Analyze package (https://github.com/bintulab/HT-recruit-Analyze) as described previously(*21*, *35*). Domains with fewer than 20 counts in both On and Off fractions of a given replicate were filtered out of downstream analyses. Hit thresholds for the Day 1 and Day 5 minCMV screen were set at 1 standard deviation above the mean of poorly expressed random control tiles. Activation domains and bifunctional domains were labeled based on annotations from minCMV activation and pEF repression screens performed in DelRosso et al., 2023(*21*).

**Measuring domain abundance with HaloTag staining**

HaloTag staining was performed using the Janelia Fluor 646 HaloTag Ligand (Promega #GA1121) as follows: ligand was resuspended in 35.5 µL DMSO to prepare a 200 µM stock solution, then diluted in a 200 nM working solution in warm RPMI. 2x$10^5$ cells were pelleted per sample and resuspended in 200 µL of working ligand solution. Samples were incubated for 15 minutes in a 37°C 5% $CO_2$ incubator, then pelleted and resuspended in 200 µL RPMI for flow cytometry analysis. Cells expressing either HaloTag or mCherry fusion proteins, as well as wild-type cells, were stained and flowed alongside experimental samples and used to compensate for mCherry - Janelia Fluor 646 bleedthrough.

**Measuring cell proliferation**

Cell growth quantification was performed using the ViaFluor 405 SE Cell Proliferation Dye (ViaFluor #30068) as follows: dye was resuspended in 20 µL DMSO to prepare a 5 µM stock solution, then diluted to a 5 µM working solution in warm RPMI. 5x$10^5$ cells were pelleted per sample and resuspended in 500 µL of working dye solution. Samples were incubated for 15 minutes in a 37°C 5% $CO_2$ incubator, then 500 µL RPMI was added to hydrolyze free dye and incubated for 5 additional minutes. Cells were pelleted and resuspended in 1 mL fresh RPMI and cultured for 4 days. Each day, 200 µL cells were analyzed by flow cytometry to detect dye dilution rate.

**Chemical inhibition of transcriptional cofactors**

500 nM dCBP-1 (MedChemExpress #HY-134582) was added to cell culture media 6 hours prior to doxycycline addition to degrade P300/CBP. 10 µM A485 (Selleck Chemicals #S8740) was added to culture media 2 hours prior to dox addition to catalytically inhibit P300/CBP. 10 µM TMP269 (Selleck Chemicals #S7324), 10 µM Tazemetostat (Selleck Chemicals #S7128), or 10 µM SETDB1-TTD-IN-1 (MedChemExpress #HY-141539) were added to culture media 24 hours prior to dox addition to inhibit Class IIa HDACs, EZH2, or SETDB1, respectively. Drug conditions were matched with a DMSO vehicle control and Citrine expression changes during dox recruitment were compared to matched no dox controls for each drug and cell line. Media and inhibitors were replenished every 24 hours as described above.

**CUT&RUN for histone modifications during domain recruitment**

CUT&RUN was performed with the CUTANA ChIC/CUT&RUN kit (EpiCypher #14-1048) according to manufacturer's instructions from manual version 4. Antibody incubations were performed overnight at 4°C and used at 1:50. Antibodies used for CUT&RUN: Anti-H3K9me3 (Abcam ab176916), Anti-H3ac (Active Motif #39139), H3K4me3 (EpiCypher 13-0041), Rabbit IgG (EpiCypher 13-0041). Libraries were prepared using the CUT&RUN Library Prep Kit (EpiCypher 14-1001/14-1002) with 5 ng of input DNA according to manufacturer's instructions. DNA was quantified using a Qubit4 fluorometer and libraries were sequenced at a targeted depth of 10M paired-end reads per sample on an Illumina NovaSeq with 2x150 cycles. A custom human genome (hg38) with the minCMV reporter gene integrated at the AAVS1 locus on chromosome 19 was created using bowtie2-build. Alignment was performed using bowtie2 and sorted

BAM files were generated using samtools. Duplicate reads were removed using Picard. Deduplicated BAM files were indexed using samtools and counts per million normalization was performed with bamCoverage to generate final bedGraph files used in downstream analysis. Processing scripts are available at https://github.com/bintulab/Spreading_Lensch_2022/tree/main/CUT%26RUN%20Analysis (*72*). Further analysis was performed using custom Matlab scripts available at https://github.com/bintulab/ORCA_Fujimori_2023/tree/main/CUTnRUN_analysis (*73*). Reporter gene reads/kb were normalized to positive and negative control genes as follows: (reporter reads/kb - negative control reads/kb)/(positive control reads/kb - negative control reads/kb).

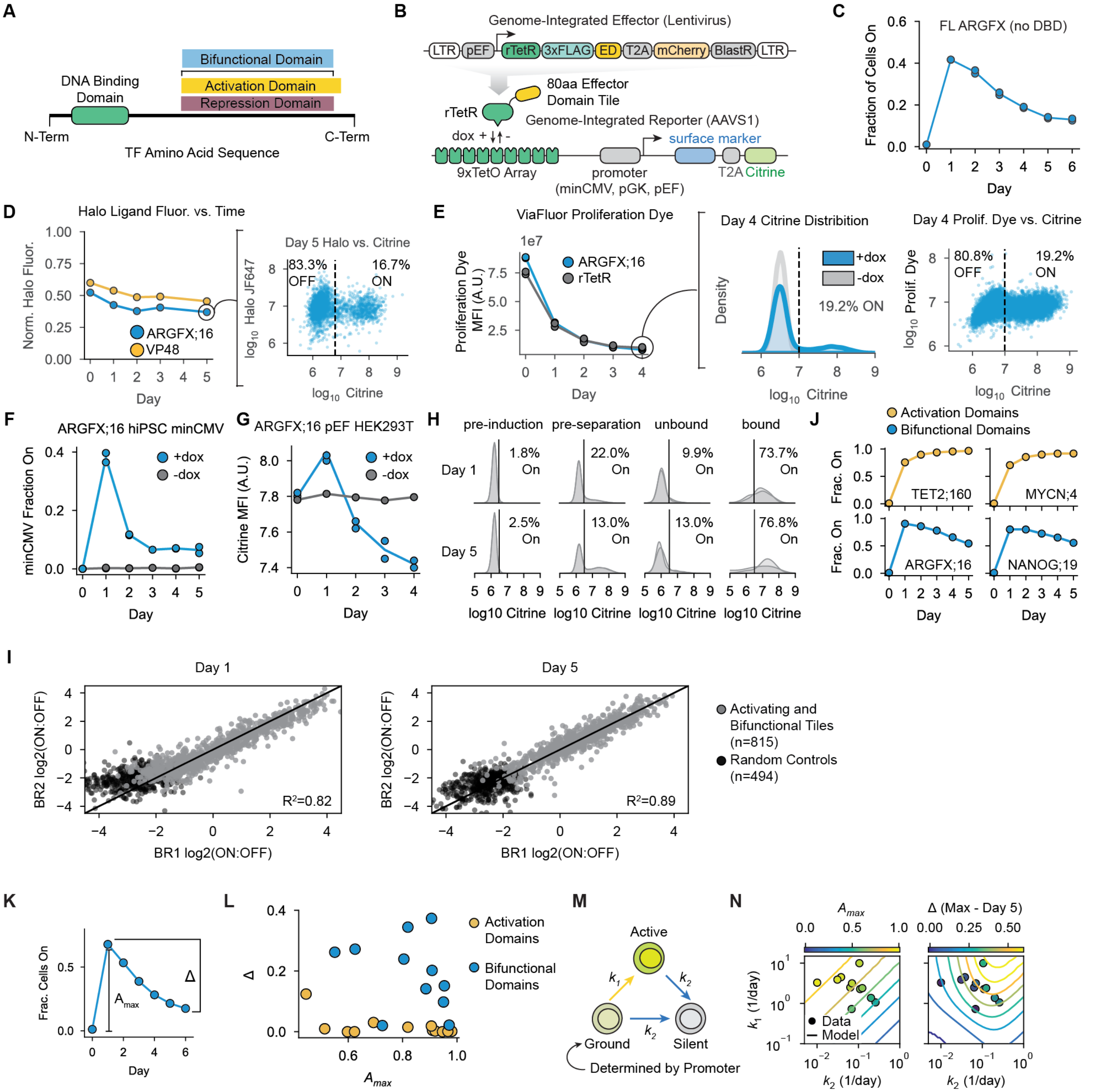


**Figure S1: Validation of bifunctional domain pulse generation and HT-Recruit dynamics measurements. A)** Schematic of an example bifunctional domain. Bifunctional domains within human TFs and chromatin regulators are defined as domains that were sufficient to activate the minimal minCMV promoter and repress the constitutively active pEF promoter in our previous study(*21*). **B)** Schematic of genome-integrated rTetR-effector domain fusion expression construct and AAVS1-integrated Citrine and surface marker reporter genes. **C)** Fraction of active cells during recruitment of full-length ARGFX without its DBD to minCMV in K562 cells. Residues 78-137 encode the ARGFX ETCHbox domain and were replaced with a GS linker. **D)** (Left) Normalized Halo Tag fluorescence during ARGFX;16 (blue) or VP48 (yellow) recruitment to minCMV reporter in K562 for 5 days. Halo Tag fluorescence is normalized to the fluorescence of Halo ligand-stained cells that do not express a Halo-tagged protein (negative control, 0) and the fluorescence of Halo-tagged rTetR alone (highly expressed positive control, 1). (Middle) Citrine fluorescence distribution on Day 5 of ARGFX;16 recruitment. (Right) Comparison of Halo Tag vs. Citrine fluorescence in single cells on Day 5 of recruitment. **E)** (Left) Mean fluorescence intensity of ViaFluor proliferation dye during 4 days of ARGFX;16 (blue) or rTetR (gray)

recruitment. (Middle) Distribution of Citrine fluorescence on Day 4 of ARGFX;16 recruitment. (Right) Comparison of ViaFluor proliferation dye fluorescence vs. Citrine fluorescence in single cells on Day 4 of recruitment. **F)** Fraction of cells On during ARGFX;16 recruitment to the minCMV promoter in hiPSCs (blue=1000 ng/mL dox, gray=0 ng/mL dox). **G)** Mean fluorescence intensity of cell population during ARGFX;16 recruitment to the pEF promoter in HEK293T cells (blue=1000 ng/mL dox, gray=0 ng/mL dox). **H)** Percentage of cells On in pre- and post-magnetic separation fractions for Day 1 and Day 5 screen. **I)** Screen reproducibility between biological replicates for Day 1 and Day 5 screen. (gray=activating and bifunctional tiles, black=random controls) (Day 1 $R^2$=0.82, Day 5 $R^2$=0.89). **J)** Fraction of cells On during activation domain or bifunctional domain recruitment to minCMV in K562 cells. **K)** Pulse of gene expression at minCMV can be summarized by shape parameters $A_{max}$ and Δ (Max fraction On - final measured fraction On). **L)** $A_{max}$ and Δ quantified for individually validated bifunctional domains (blue) and activation domains (yellow). **M)** Minimal kinetic model of bifunctional regulation based on stochastic transitions between ground, active, and silent states. Yellow arrows indicate transitions towards an active state, blue arrows indicate transitions towards a silent state. **N)** Contour plot showing minimal model predictions for $A_{max}$ and Δ for varying values of $k_1$ and $k_2$. Circles represent fit values of $k_1$ and $k_2$ from 11 individually validated pulse-generating domains from 9 unique TFs.

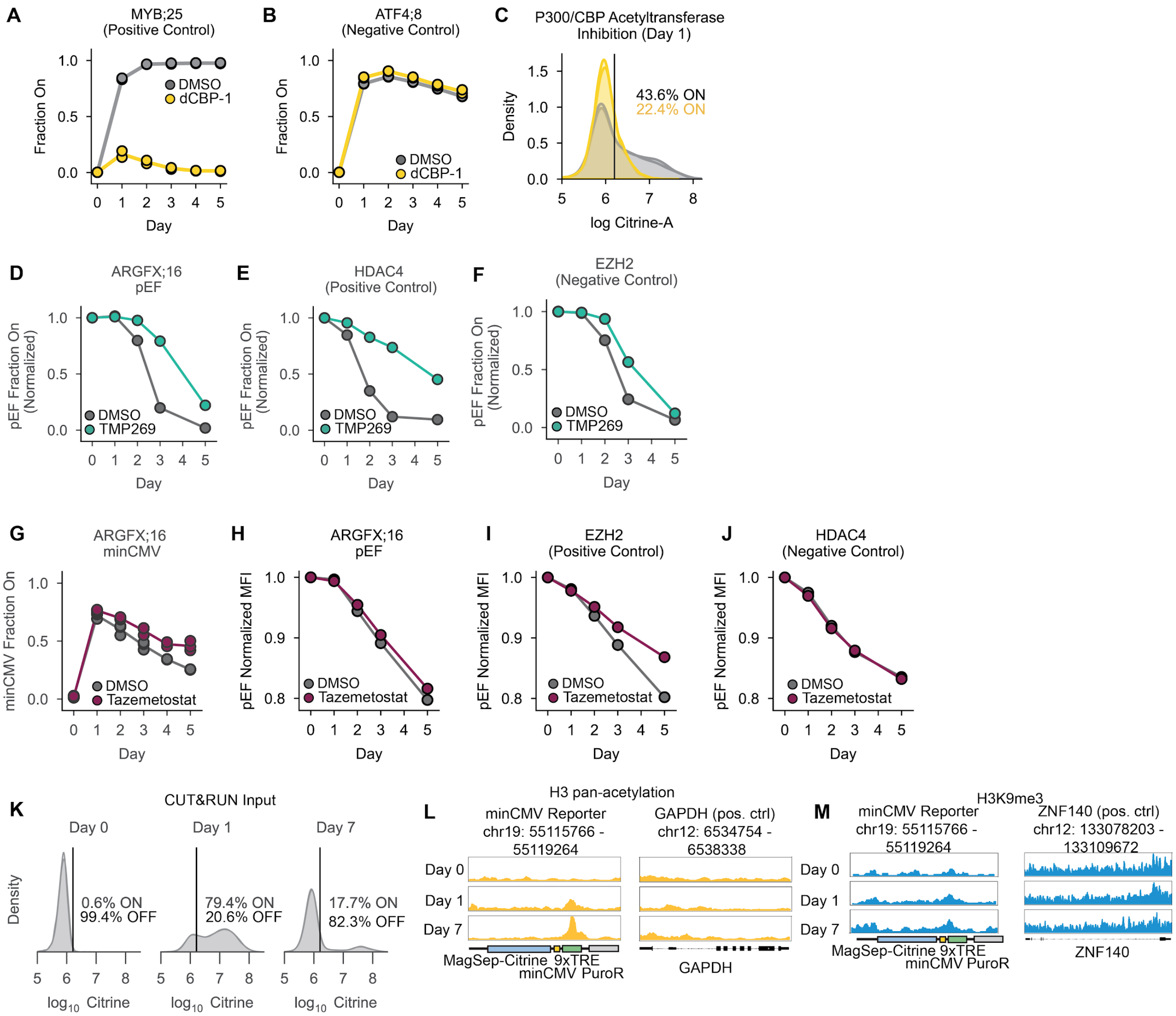


**Figure S2: Testing dependence of pulse dynamics on transcriptional cofactors and histone modifications. A)** MYB;25 known P300 binder (positive control) recruited to minCMV in DMSO or 10 µM dCBP-1 media. **B)** ATF4;8 P300-independent activator (negative control) recruited to minCMV in DMSO or 10 µM dCBP-1 media. **C)** Distribution of Citrine expression after 1 day of ARGFX;16 recruitment with 10 µM A485 P300/CBP acetyltransferase small molecule inhibitor (yellow) or DMSO (gray). **D)** Fraction of cells On during ARGFX;16 recruitment to pEF in DMSO (vehicle control, gray) or 10 µM TMP269 (Class IIa HDAC inhibitor, green) media normalized to 0 ng/mL dox control. **E)** HDAC4 catalytic domain (positive control) recruited to pEF in DMSO or 10 µM TMP269 media. **F)** EZH2 catalytic domain (negative control) recruited to pEF in DMSO or 10 µM TMP269 media. **G)** ARGFX;16 recruited to minCMV in DMSO (vehicle control, gray) or 10 µM Tazemetostat (EZH2 inhibitor, maroon) media. **H)** Mean fluorescence intensity (MFI) during ARGFX;16 recruitment to pEF in DMSO or 10 µM Tazemetostat media normalized to 0 ng/mL dox control. **I)** MFI during EZH2 (positive control) recruitment to pEF in DMSO or 10 µM Tazemetostat media. **J)** MFI during HDAC4 (negative control) recruitment to pEF in DMSO or 10 µM Tazemetostat media. **K)** Citrine fluorescence distribution of CUT&RUN input cell populations. **L)** H3 pan-acetylation genome browser tracks at minCMV reporter or GAPDH (H3 pan-acetylation positive control gene) after 0, 1, or 7 days of ARGFX;16 recruitment. **M)** H3K9me3 genome browser tracks at minCMGV reporter or ZNF140 (H3K9me3 positive control gene) after 0, 1, or 7 days of ARGFX;16 recruitment.

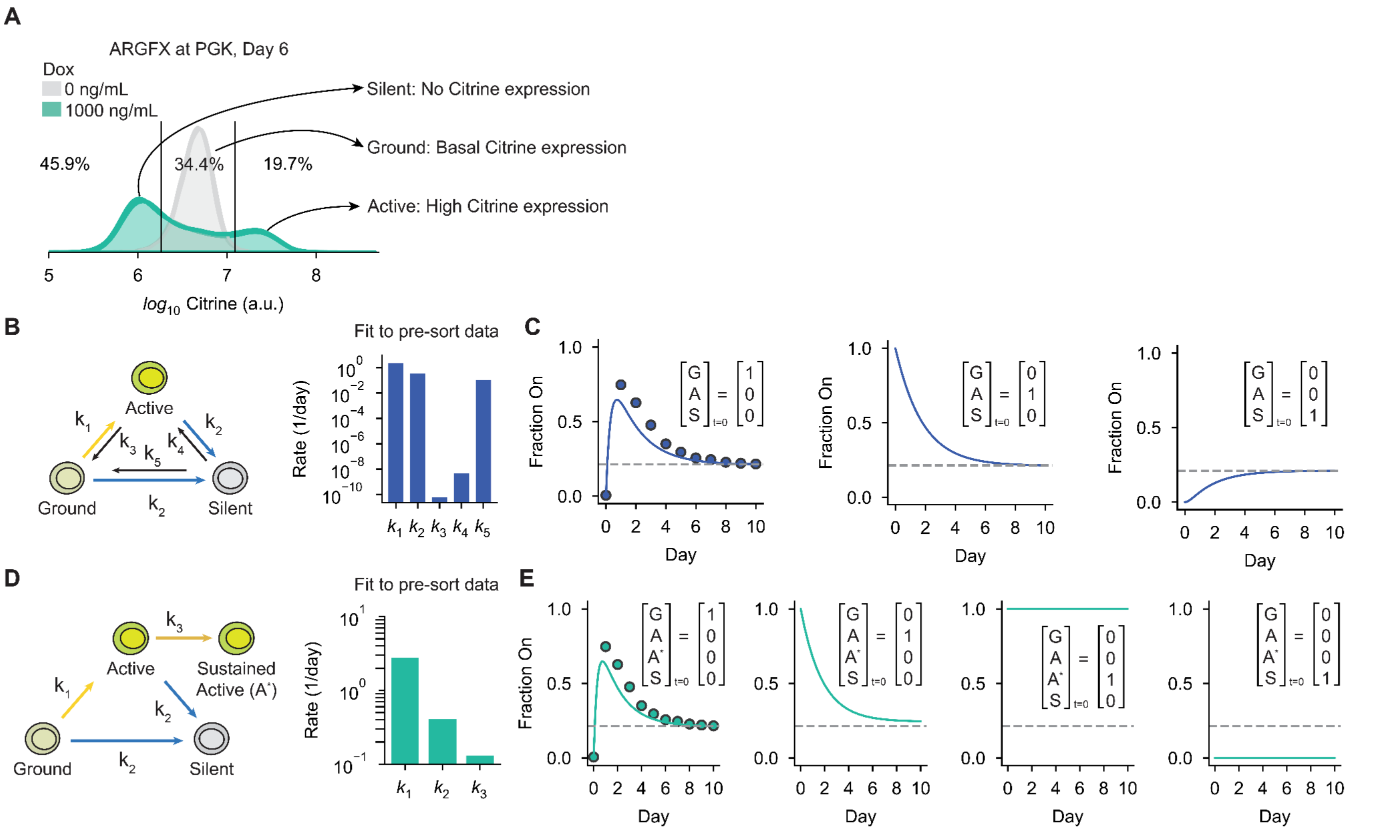


**Figure S3: Developing and distinguishing between various state-based kinetic models of bifunctional gene regulation. A)** Observation of three, discrete gene expression states created by ARGFX;16 recruited to the PGK promoter for 6 days. **B)** (Left) Enumeration of rates in the Dynamic Equilibrium model (Figure 3B, left) and (right) parameter values obtained by fitting this model to the fraction of Citrine expressing cells over 10 days of ARGFX;16 recruitment (Figure 3A). **C)** Predicted gene expression dynamics from the Dynamic Equilibrium model using the rate parameters shown in C for populations of cells starting in different gene expression states. G = ground state, A = active state, S = silent state. Dots are data used for fitting. **D)** (Left) Enumeration of rates in the Divergent States model (Figure 3B, right) and (right) parameter values obtained by fitting this model to the fraction of Citrine expressing cells over 10 days of ARGFX;16 recruitment (Figure 3A). Cells in both the active and sustained active states are assumed to express Citrine. **E)** Predicted gene expression dynamics from the Divergent States model using the rate parameters shown in E for populations of cells starting in different gene expression states. G = ground state, A = active state, S = silent state, A* = sustained active state. Data used to fit the model shown as dots. Model predictions in Figure 3D are made by *in silico* sorting Citrine expressing and silent cells using the fits in C and E and propagating gene expression dynamics forward in time.

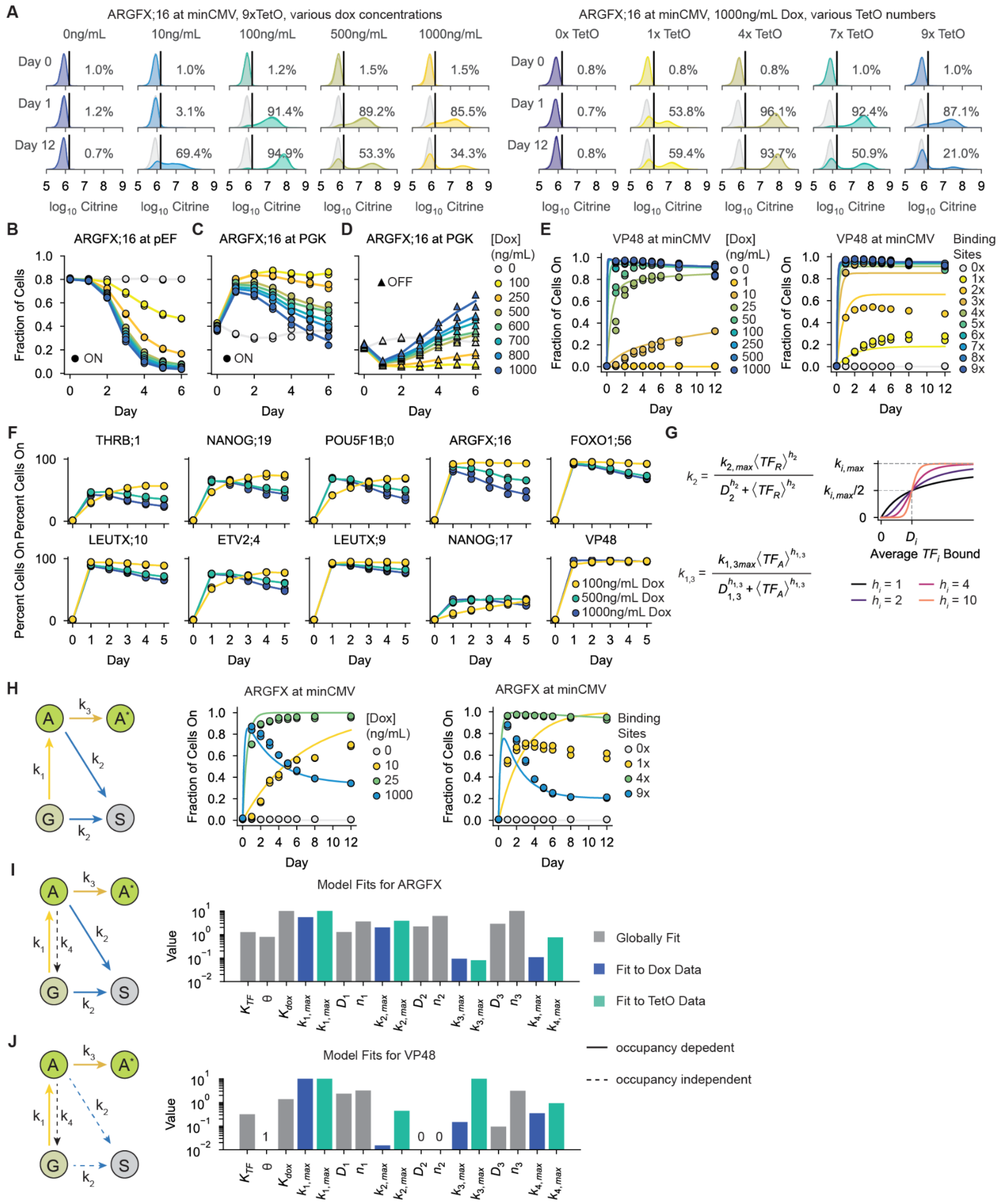


**Figure S4: Measuring and modeling the occupancy-dependence of activation and repression by bifunctional domains. A)** (Left) Citrine fluorescence distributions during ARGFX;16 recruitment at minCMV, 9 TetO sites, for a range of dox concentrations. (Right) Citrine fluorescence distributions during ARGFX;16 recruitment at minCMV using 1000ng/mL dox with different TetO copy numbers in the reporter gene. **B)** Fraction of cells expressing Citrine during ARGFX;16 recruitment to the pEF promoter at several dox concentrations. **C)** Fraction of active cells during ARGFX;16

recruitment to the PGK promoter at several dox concentrations. Cells expressing Citrine at least one standard deviation above the mean of the -dox PGK Citrine fluorescence distribution are considered active. **D)** Fraction of silenced cells during ARGFX;16 recruitment to the PGK promoter at several dox concentrations. Cells expressing Citrine at least one standard deviation below the mean of the -dox PGK Citrine fluorescence distribution are considered silenced. **E)** (Left) Fraction of cells expressing citrine during VP48 recruitment to the minCMV promoter with varying dox, 9x TetO sites. (Right) Fraction of cells expressing citrine during VP48 recruitment to the minCMV promoter with various TetO copy numbers, 1000ng/mL dox. Dots are measurements, lines are model fits from I. **F)** Fraction of cells expressing citrine over time when recruiting 9 pulse generating bifunctional domains to the minCMV promoter at three dox concentrations. **G)** Intuition behind the calculation of transition rates between gene expression states from TF occupancy. Transition rates are assumed to be Hill functions of TF occupancy. **H)** Predicted gene expression dynamics when fitting the Divergent States model (no transition rate from the active to ground state) to the data in Figure 4A-B. For clarity, four dox concentrations and numbers of binding sites are shown, but the model was fit to all data. **I)** Resulting parameters from fitting the Divergent States model (with an active to ground transition rate) to expression data during ARGFX;16 recruitment to minCMV (Figure 4A-B). **J)** Resulting parameters from fitting a modified Divergent States model (an occupancy-independent silencing rate) to expression data during VP48 recruitment to minCMV (Figure S4E). Maximum transition rates were individually fit for the dox titration and binding site perturbation experiments, all other parameters were globally fit. For VP48, all TF was assumed to be interacting with the coactivator and the rate of silencing was assumed to be independent of TF occupancy to represent background silencing.

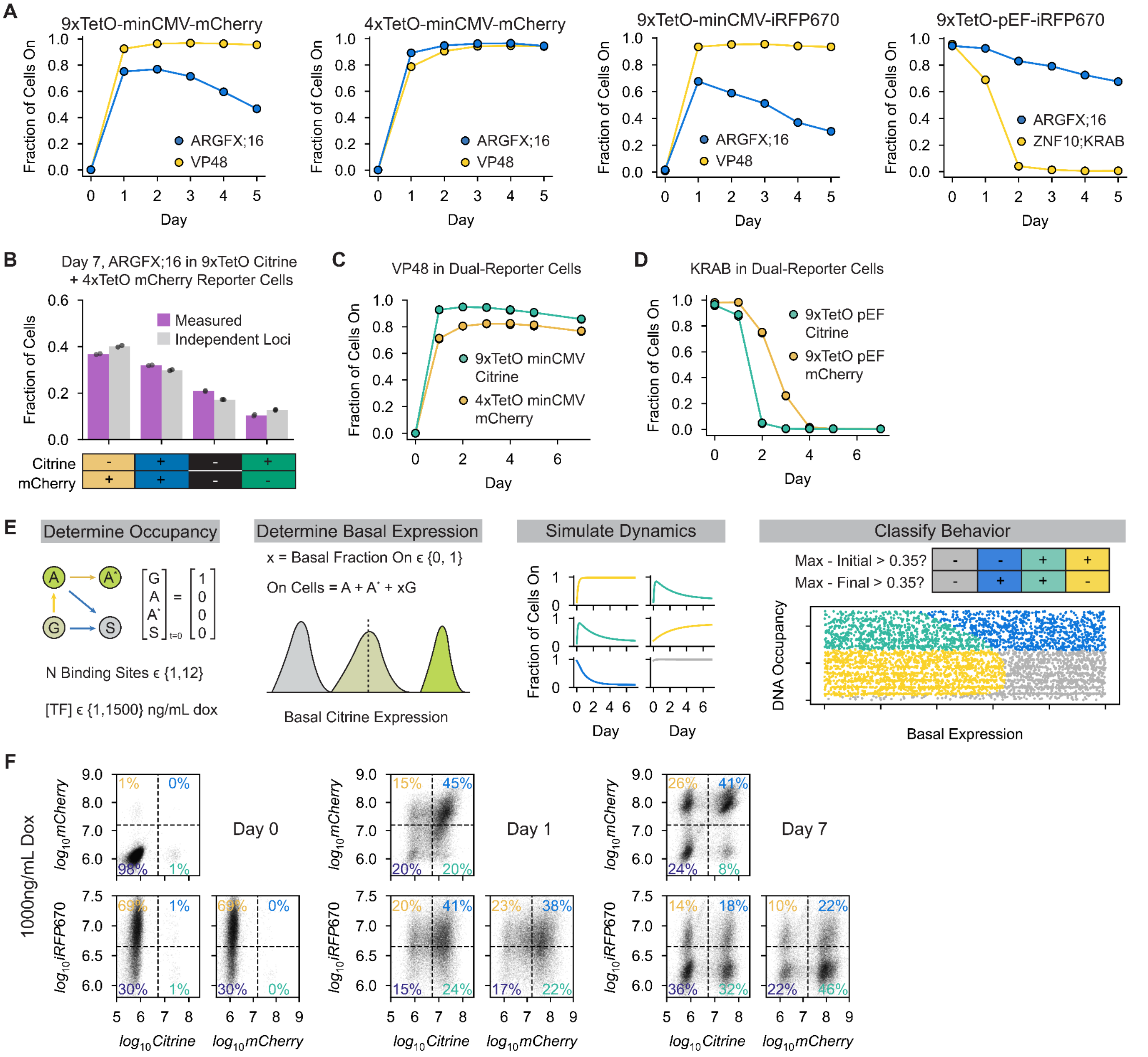


**Figure S5: Exploring the behavior space of bifunctional domains in individual cells. A)** Fraction of cells expressing various reporter genes during ARGFX;16 and VP48 recruitment in single-reporter cells. **B)** Comparison of measured fraction of cells (magenta) in each quadrant and the expected fraction of cells (gray) in each quadrant under the assumption that each reporter is regulated completely independently of the other after 7 days in ARGFX;16 recruitment in dual reporter cells (Figure 5B). The "independent loci" expectation is the product of the probability of each gene being on or Off based on population-wide measurements (Figure 5C). Small deviations from perfect independence are likely caused by background silencing of the recruiter construct resulting in a slight enrichment of double-negative cells. **C)** Fraction of cells expressing Citrine when recruiting VP48 in dual-reporter cell line with one reporter containing 9 TetO sites upstream of minCMV and the other containing 4 binding sites upstream of minCMV, 1000ng/mL dox. **D)** Fraction of cells expressing Citrine when recruiting KRAB in dual-reporter cell line with each reporter containing 9 TetO sites upstream of pEF, 1000ng/mL dox. **E)** Procedure for generating the extended behavior space of bifunctional domains shown in Figure 5G. Gene expression dynamics are simulated over 12 days. **F)** Single-cell scatterplots of Citrine, mCherry, and miRFP670nano3 expression during ARGFX;16 recruitment in the triple-reporter cell line at 1000ng/mL dox. Dashed lines are the gates used to calculate the fraction of cells expressing each reporter in Figure 5H-K.
